# The ß-catenin-target Fascin-1, altering hepatocyte differentiation, is a new marker of immature cells in hepatoblastomas

**DOI:** 10.1101/2021.04.21.440735

**Authors:** Caroline Gest, Sandra Sena, Véronique Neaud, Robin Loesch, Nathalie Dugot-Senant, Lisa Paysan, Léo Piquet, Terezinha Robbe, Nathalie Allain, Doulaye Dembele, Catherine Guettier, Paulette Bioulac-Sage, Brigitte Le Bail, Christophe F. Grosset, Frédéric Saltel, Valérie Lagrée, Sabine Colnot, Violaine Moreau

## Abstract

**BACKGROUND & AIMS:** ß-catenin is a well-known effector of the Wnt pathway and a key player in cadherin-mediated cell adhesion. Oncogenic mutations of ß-catenin are highly frequent in pediatric liver primary tumors. Those mutations are mostly heterozygous allowing the co-expression of wild-type (WT) and mutated ß-catenins in tumor cells. We investigated the interplay between WT and mutated ß-catenins in liver tumor cells, and searched for new actors of the ß-catenin pathway.

**METHODS:** Using an RNAi strategy in ß-catenin-mutated hepatoblastoma (HB) cells, we dissociated the structural and transcriptional activities of β-catenin, carried mainly by, respectively, WT and mutated proteins. Their impact was characterized using transcriptomic and functional analyses. We studied mice that develop liver tumors upon activation of ß-catenin in hepatocytes (APC^KO^ and ß-catenin^Δexon3^ mice). We made use of transcriptomic data from mouse and human HB specimens and analyzed samples by immunohistochemistry.

**RESULTS:** We highlighted an antagonist role of WT and mutated ß-catenins on hepatocyte differentiation as attested by alteration of hepatocyte markers expression and bile canaliculi formation. We characterized Fascin-1 as a target of ß-catenin involved in hepatocyte differentiation. Using mouse models that allow the formation of two phenotypically distinct tumors (differentiated or undifferentiated), we found that Fascin-1 expression is higher in undifferentiated tumors. Finally, we found that Fascin-1 is a specific marker of the embryonal component in human HBs.

**CONCLUSIONS:** In mice and human, Fascin-1 expression is linked to loss of differentiation and polarity of hepatocytes. Thus, we highlighted Fascin-1 as a new player in the modulation of hepatocyte differentiation associated to ß-catenin pathway alteration in the liver.

**Data Transparency Statement:** study materials will be made available to other researchers upon request.

## INTRODUCTION

ß-catenin is an evolutionary conserved protein that plays a dual role in cells. It is the key effector of the canonical Wnt pathway, acting as a transcriptional cofactor in complex with lymphoid enhancer factor/T-cell factor (LEF/TCF)^1^. In addition, ß-catenin plays a central role in cadherin-mediated cell adhesion. In epithelial cells, in the absence of Wnt signaling, ß-catenin is associated with E-cadherin at cell-cell junctions, and β-catenin is maintained at low cytoplasmic levels through its destruction complex of Axin, adenomatosis polyposis coli (APC), and glycogen synthase 3ß kinase (GSK-3ß). GSK-3ß phosphorylates ß-catenin causing its degradation by the proteasome. Upon Wnt signal, the destruction complex is disrupted and cytoplasmic stabilized ß-catenin translocates in the nucleus where it drives the transcription of target genes. Thus, ß-catenin is endowed with two main functions, a structural function at cell-cell adhesion and a transcriptional function in the nucleus. Imbalance between signaling properties of ß-catenin may lead to deregulated cell growth, adhesion and migration resulting in disease such as tumor development and metastasis. However, given the dual function of ß-catenin, it is difficult to distinguish which function, structural versus transcriptional, of ß-catenin, is precisely involved in cellular processes.

In the liver, the Wnt/ß-catenin pathway plays important roles regulating embryonic and postnatal development, zonation, metabolism and regeneration^2^. This pathway is also strongly involved in the hepatocarcinogenesis. Its aberrant activation, due to mutations in the *CTNNB1* gene encoding ß-catenin or in components of the degradation complex, such as *AXIN* and *APC*, are detected in primary liver cancers. ß-catenin gene alterations are identified in up to 80% of human hepatoblastomas (HBs), the primary hepatic malignancy in children, and in 30 to 40% of human hepatocellular carcinomas (HCCs)^3, 4^. Moreover, whereas *APC* loss-of function mutations remain rare in HCCs, they are more frequent in HBs, associated to familial or sporadic cases^5^. Even if HBs remain poorly studied and complex tumors with various histologic components (epithelial, mesenchymal, fetal, and embryonal) within the same tumor^6^, two subtypes were described based on molecular analyses; the fetal C1 subtype with favorable outcome shows enhanced membranous staining and cytoplasmic accumulation of ß-catenin with occasional nuclear localization, whereas the proliferative poorly differentiated C2 subtype is characterized by an intense nuclear staining of ß-catenin^3^. The C2 subtype was recently split into subgroups, C2A and C2B, with C2A subgroup containing more proliferative tumors^7^. The majority of ß-catenin mutations affect exon3 at sites of phosphorylation by GSK3ß, avoiding its degradation and so constitutively activating the Wnt pathway. Interstitial deletions in the third exon of the ß-catenin gene are highly prevalent in HBs, while point mutations are more common in HCCs^8^. Interestingly, most of mutations in exon3 are monoallelic mutations, leaving a wild-type (WT), non-mutated allele in tumor cells. We thus attempt to address the interplay between the WT and the mutated ß-catenins in liver tumor cells.

To do so, we made use of the human HB HepG2 cell line. HepG2 cells have an heterozygous deletion of 348 nucleotides in exon3 of the *CTNNB1* gene, resulting in an abundant truncated form of ß-catenin and a lower amount of WT ß-catenin^4^. The large deletion (amino acid residues 25–140) removes the GSK-3ß phosphorylation sites and the binding site for α-catenin, the ß-catenin partner in the E-cadherin-mediated cell adhesion. Thus, both WT and mutated (Δaa25-140) forms of ß-catenin co-exist in these cells and the interplay between both has never been explored. In addition, HepG2 cells exhibit a good degree of differentiation and show most cellular features of normal human hepatocytes such as bile canaliculi (BC) formation^9^. We thus designed an RNA interference approach to specifically knockdown WT and/or mutated ß-catenin and address their reciprocal role in hepatocyte differentiation. Using this model, we dissociated the structural and transcriptional activities of β-catenin, carried mainly by, respectively, the WT and the mutated proteins. Moreover, we found that whereas knockdown (KD) of WT ß-catenin induced a loss of BC, KD of mutated ß-catenin lead to an increase of their number and size, suggesting that both play antagonistic role in tumor hepatic cell differentiation. Moreover, molecular characterization revealed that *FSCN1* (fascin-1) is a target of ß-catenin involved in the differentiation state of hepatocytes. We further found that fascin-1 expression is associated to undifferentiated ß-catenin mutated tumors in mice, which are closed to human HBs. Using human samples, we found that fascin-1 is specifically expressed in the embryonal component of HBs. Thus, we described *FSCN1* as a ß-catenin target gene associated with hepatic tumors of poor outcome, such as poorly differentiated HBs.

## MATERIALS AND METHODS

### Cell culture

Human HB cell line, HepG2, and human HCC cell lines, Hep3B, Huh7 and SNU398, were purchased from American Type Culture Collection. HB cell line Huh6 was generously provided by C. Perret (Paris, France). All cell lines were cultured as previously described^10, 11^.

### Transfection

DNA and siRNA transfections were realized as described previously^11^. pEGFP-Fascin-1 vector was a generous gift from Dr D. Vignjevic (Paris, France). SiRNA were purchased from Eurofins genomics. Sißcat WT targets human ß-catenin mRNA at 5’-GTAGCTGATATTGATGGACAG-3’, sißcat both at 5’-ACCAGTTGTGGTTAAGCTCTT-3’ and sißcat mut at 5’-TGTTAGTCACAACTATCAAGA-3’. SiFascin-1#1 targets human Fascin-1 mRNA at 5’-CAGCTGCTACTTTGACATCGA-3’, siFascin-1#2 at 5’-CAAAGACTCCACAGGCAAA-3’, siFascin-1#3 and siFascin-1#4 were purchased at Ambion. The AllStars negative-control siRNA from Qiagen was used as control siRNA. For KD, a reverse transfection was performed on day 1, a second forward transfection on day 3 and experiments were performed on day 5.

### Luciferase reporter assays

Assays were performed as previously described^11^. pGL4-TOP reporter with TCF responsive elements to quantify ß-catenin transcriptional activity was a generous gift from Pr. H. Clevers (Utrecht, The Netherlands). The pmFascin-Luciferase vector containing luciferase under fascin promoter was a generous gift from Dr D. Vignjevic (Paris, France).

### Western blot

Cells were scraped off on ice and homogenized in RIPA buffer (150 mM NaCl, 0.1% Triton X**-**100, 0.5% sodium deoxycholate, 0.1% sodium dodecyl sulfate, and 50 mM Tris**-**HCl pH 8.0) containing protease and phosphatase inhibitors cocktail (Thermo Scientific). Cell lysates were cleared of cellular debris and nuclei by a 16,000 × g centrifugation step for 10 min. Lysates were analyzed using a western-blot protocol described previously^11^. The antibodies used are listed in Table S1.

### Quantitative real-time PCR

RNA was collected from cultured cells using the Trizol reagent (Invitrogen), according to manufacturer’s protocol. Reverse transcription and SYBR® Green-based real-time PCR were performed as described previously^11^. Gene expression results were first normalized to internal control r18S. Relative levels of expression were calculated using the comparative (2^-ΔΔCT^) method. All primers used for qRT-PCR experiments are listed in Table S2.

### Immunofluorescence

Glass coverslip-plated cells were prepared for immunofluorescence microscopy and imaged as previously described^12^. The antibodies used are listed in Table S1. F-actin was stained using fluorescent phalloidin (1/250) (Molecular Probes). To detect functional BC, cells were incubated in PBS with 5-carboxyfluorescein diacetate (CFDA) (Sigma) at a final concentration of 0.5 µM for 30 min at 37°C to allow its internalization into the lumen. Cells were washed three times with PBS and CFDA positive BC were counted as functional under a fluorescence microscope. Stimulated emission depletion (STED) microscopy was used to image microvilli of BC, by labelling actin with phalloidin-Atto647N (Thermo Fischer Scientific). Sample was imaged with an inverted Leica SP8 STED microscope equipped with an oil immersion objective (Plan Apo 100X NA 1.4), white light laser (WLL2) and internal hybrid detectors. BC features were quantified using image J on STED images. Quantification of BC was assessed in three independent experiments in which at least 100 cells were counted.

### Cell growth assay

For 2D assay, cells were seeded at 5000 cells/well in 96-well plates and grown for 1 to 6 days. Each assay was performed in five replicates. For indicated time points, cells were fixed with 50% trichloroacetic acid at 4°C for 1 hour, and processed using the SulfoRhodamine B assay kit (Sigma).

### Mouse samples

We collected tumoral and non tumoral livers from mouse transgenic models with hepatic β-catenin activation. All animal procedures were approved by the ethical committee of Université de Paris according to the French government regulation. The *Apc*^fs-ex15^ and β-catenin^Δex3^ mouse tumors were obtained from compound *Apc*^flox/flox^/TTR-Cre^Tam^ and β-catenin^ex3-flox/flox^ respectively, injected with 0.75 mg Tamoxifen or with 5 x 10^8^ ip Cre-expressing adenovirus, as previously described^13–15^. All the mice were maintained at the animal facility with standard diet and housing. They were followed by ultrasonography every month until tumor detection, thereafter ultrasound imaging was continued every 2 weeks. Table S3 described the cohort of mice used for this study.

### Patient samples

Liver tissues were immediately frozen in Isopentane with Snapfrost and stored at −80°C until used for molecular studies. Samples were obtained from the Centre de Ressources Biologiques (CRB)-Paris-Sud (BRIF N°BB-0033-00089) with written informed consent, and the study protocol was approved by the French Government and the ethics committees of HEPATOBIO (HEPATOBIO project: CPP N°CO-15-003; CNIL N°915640). Liver samples were clinically, histologically, and genetically characterized (Table S4). Among the 20 cases, 12 were classified as C1, 3 as C2A and 5 as C2B in a C. Grosset’s previous study^7^.

### Immunohistochemistry

Specimens were fixed in buffered formaldehyde. The 2.5 µm thick sections were dewaxed and rehydrated and antigen retrieval was performed in a Tris-EDTA buffer (pH9 solution for 20 min). All staining procedures were performed in an automated autostainer (Dako-Agilent Clara, United States) using standard reagents provided by the manufacturer. Endogenous peroxidase was inhibited with 3% H_2_O_2_ in H_2_O, and non-specific sites were saturated with 20% goat serum in TBS-Tween. The sections were incubated with the anti-Fascin-1 monoclonal antibody at 2 µg/mL for 45 min at room temperature. EnVision Flex/HRP was used for signal amplification. 3,3’-Diamino-benzidine development was used for detecting primary antibodies. The slides were counterstained with hematoxylin, dehydrated and mounted.

### Statistical tests

Data were reported as the mean ± SEM of at least three experiments. Statistical significance (P < 0.05 or less) was determined using a Student’s *t-*test or analysis of variance (ANOVA) as appropriate and performed with GraphPad Prism software. *P* values are indicated as such: * *P* < 0.05; ** *P* < 0.01; *** *P* < 0.001; **** *P* <0.0001; *ns*, non significant.

## RESULTS

### The dual ß-catenin knocked-down HepG2 model

The N-terminus part of ß-catenin contains the GSK-3ß phosphorylation sites that are crucial for the regulation of its turnover. This region encoded by the exon3 of ß-catenin is deleted on one allele of the *CTNNB1* gene in HepG2 cells^4^. Thus, a high amount of a truncated form of 76kDa is co-expressed with a 92kDa full-length ß-catenin (Fig. 1). We designed small interfering RNAs (siRNAs), named “sißcat-WT”, “sißcat-mut” and “sißcat-both” to target, respectively and specifically, either the WT, the mutated form of ß-catenin, or both (Fig. 1A). All three siRNAs efficiently knocked-down the protein expression of their targeted form of ß-catenin (Fig. 1B and S1A). At the mRNA level, we found that sißcat-WT and sißcat-mut decreased the amount of ß-catenin transcripts by two fold and that, as expected, sißcat-both led to almost a full extinction of ß-catenin in HepG2 cells (Fig. 1C).

**Figure 1:**
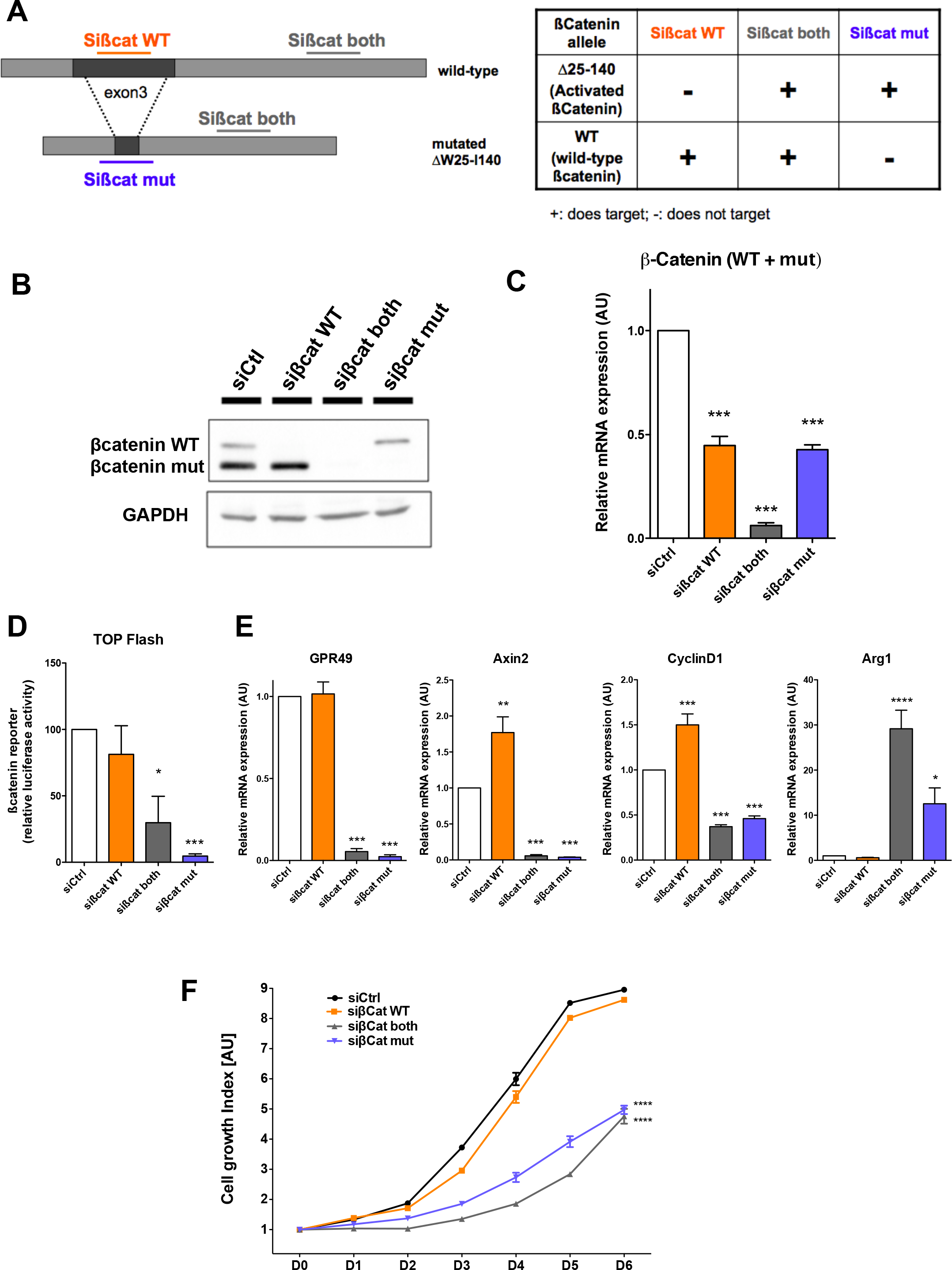
The HepG2 dual ß-catenin KD model. (A) Schema showing the different siRNAs used to KD wild-type (WT), mutated or both ß-catenins on the two alleles of *CTNNB1* gene in HepG2 cells. (B-E) HepG2 cells were transfected with indicated siRNAs. (B) Protein extracts were analyzed using anti-ß-catenin and GAPDH antibodies. Note that due to the W25-I140 deletion, mutated ß-catenin (about 76 kDa) migrates faster and is more abundant (because, more stable) than the WT ß-catenin (about 92 kDa). (C) ß-catenin mRNA expression was analyzed by qRT-PCR. (D) Promoter activity was evaluated by luciferase reporter assays in HepG2 cells transfected with TCF responsive reporter. Shown is the mean relative luciferase activity, normalized to Renilla luciferase and compared to siRNA control transfected cells. (E) mRNA levels of indicated genes were analyzed by qRT-PCR. *LGR5* (GPR49), Axin2 and *CCND1* (cyclinD1) are positive transcriptional targets of ß-catenin. *ARG1* encoding arginase is a negative transcriptional target of ß-catenin. Shown is the relative mRNA level compared to control transfected cells. (C-E) Each graph shows the quantification of three independent experiments. Error bars indicate s.e.m (n=3), *P* values from one way ANOVA. (F) HepG2 cells transfected with indicated siRNAs were seeded in 96-well plates and cells were fixed and total biomass, reflecting the number of cells, was assayed every day. Each time point was performed in triplicates. Error bars indicate s.e.m (n=3). *P* value from one way ANOVA.

We then characterized this model by analyzing the transcriptional activity of ß-catenin upon silencing of both alleles. Using the TOP-flash reporter system, we found that whereas silencing the WT allele did not impact TCF/LEF reporter activity, silencing the expression of the mutated allele or both alleles strongly inhibited it (Fig. 1D). We further showed that expression of positive targets of ß-catenin, such as GPR49, Axin2 and cyclinD1 were strongly inhibited upon treatment with sißcat-mut and sißcat-both (Fig. 1E). Consistently, the reverse result was obtained for negative targets of ß-catenin such as Arg1 (Fig. 1E). Thus, the data suggest that, in HepG2 cells, the TCF-dependent transcriptional activity of ß-catenin is mainly carried by the mutated form of ß-catenin.

According to this impact on ß-catenin targets, we found that HepG2 cell growth was highly dependent on mutated ß-catenin expression, whereas it was insensitive to the KD of the WT form of ß-catenin (Fig. 1F). This data obtained in 2D culture, were also confirmed on spheroids (Fig. S1B), showing that the mutated ß-catenin is required for HepG2 cell growth in a 2D and 3D environment. This alteration of cell growth correlated with the down-expression of cyclinD1 upon mutated ß-catenin KD (Fig. 1E). Thus, these results confirm that the mutated form of ß-catenin acts as an oncogene independently of the WT ß-catenin, and consistent with the notion of oncogene addiction, this allele is strictly required for HepG2 cell growth.

The large deletion (residues 25–140) present in HepG2 cells also removes α-catenin binding site of ß-catenin, involved in the formation of E-cadherin-based adherens junctions. We thus found that the silencing of the mutated allele of ß-catenin did not impact E-cadherin localization at cell-cell junctions. Interestingly, in an opposite manner, WT ß-catenin KD strongly affected E-cadherin engagement at adherens junctions in HepG2 cells (Fig. 2A). In cells lacking the expression of both ß-catenins, E-cadherin staining appears strongly affected with a more intracellular localization. Thus, these results suggested that the structural role of ß-catenin is altered only upon WT ß-catenin KD. Thus, this dual ß-catenin KD HepG2 model allows the uncoupling of the two functions of ß-catenin in the same cellular background: membrane/structural activity mediated by the degradable WT ß-catenin and the nuclear/transcriptional activity mediated by the mutated β-catenin. This conclusion is further supported by ß-catenin localization (Fig. 2B). In control HepG2 cells, ß-catenin localized at cell-cell junctions, in the cytoplasm and in nuclei, stainings which are lost when cells are transfected with sißcat-both. In cells transiently transfected with sißcat-mut, remaining ß-catenin, i.e. WT ß-catenin, is enriched at cell-cell junctions, but failed to localize in the cytoplasm and in nuclei. At the opposite, in cells transfected with sißcat-WT, remaining ß-catenin, i.e. mutated ß-catenin, is still cytosolic and less at the membrane (Fig. 2B). These results suggest that, in HepG2 cells, WT β-catenin is mainly involved in adherent junctions whereas mutated β-catenin is preferentially involved in the regulation of gene expression. Based on these results, we believe that this dual ß-catenin KD HepG2 model is suitable to address independently the structural and the transcriptional functions of ß-catenin.

**Figure 2:**
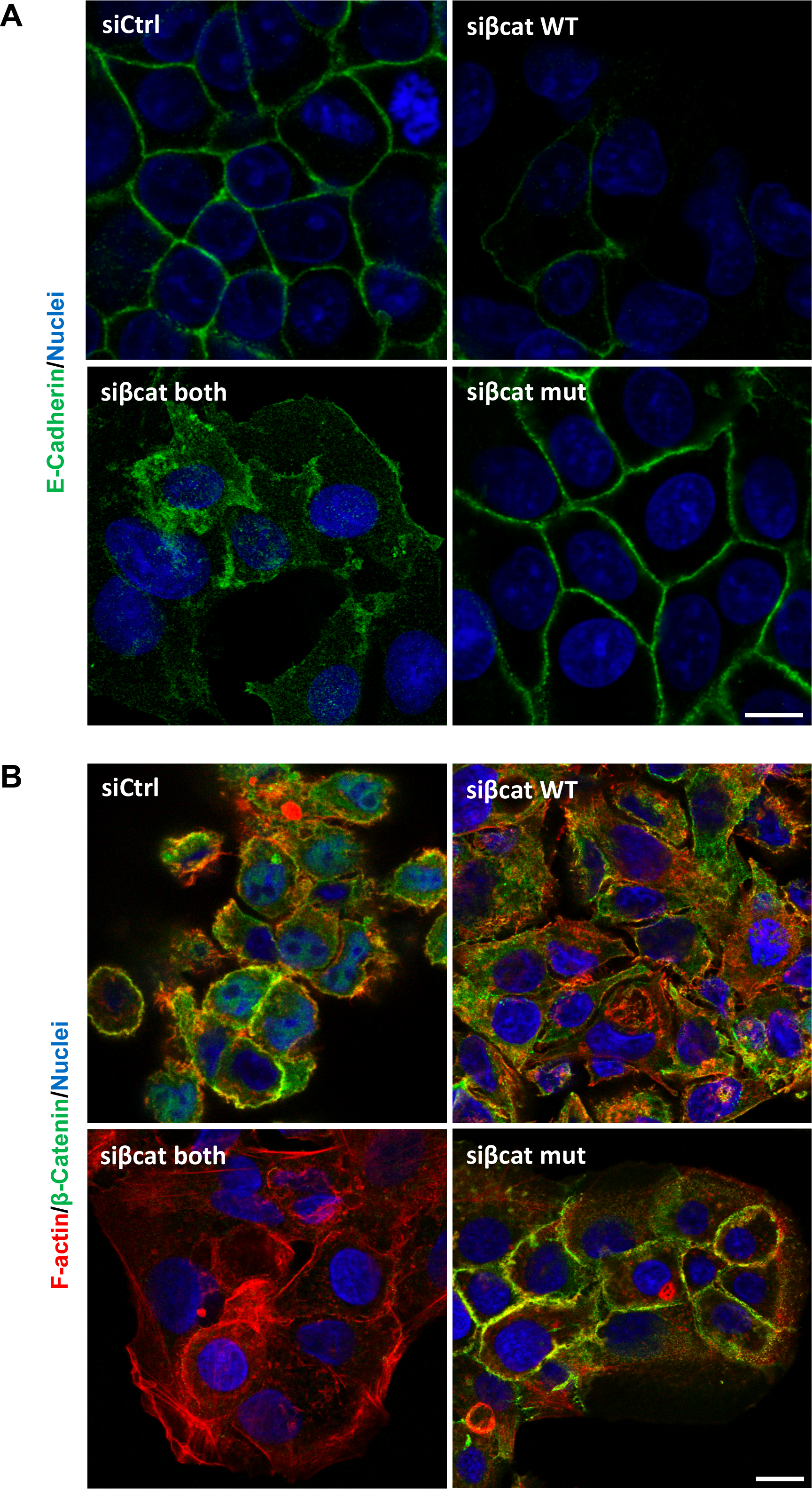
HepG2 cells transfected with indicated siRNAs were fixed and stained with (A) anti-E-Cadherin antibodies (green) and Hoescht (blue), or (B) phalloidin (red), anti-ß-catenin antibodies (green) and Hoescht (blue). Scale bar: 10 µm.

### WT and mutated ß-catenin have distinct gene expression patterns

To identify the impact of the silencing of each ß-catenin allele on gene expression, we performed a transcriptional analysis of HepG2 KD cells (Tables S5-7). To validate the data, we first checked the expression of ß-catenin and known-ß-catenin targets (Fig. S2A); they were altered in a similar way than previously observed by qRT-PCR (Fig. 1C and 1E). Global analysis demonstrated that WT ß-catenin KD cells and mutated ß-catenin KD cells have distinct gene expression patterns, and silencing of both alleles led to a gene expression pattern closer to the mutated than the WT ß-catenin KD, suggesting a dominant impact of the oncogenic ß-catenin on gene expression in HepG2 cells (Fig. S2B). Venn diagram revealed that only a small proportion of genes are altered in common upon silencing of WT or mutated ß-Catenin (Fig. S2C). We further used FuncAssociate 3.0 to analyze alterations in pathway and biological functions (Table S8). We found that removal of the WT ß-catenin led to a decrease in expression of genes involved in metabolic processes. It was striking to observe an opposite behavior upon removal of the oncogenic ß-catenin. We also found a cell cycle and a TGF-ß signature for genes that were down-regulated upon silencing of both or of the mutated allele. As metabolic functions are key features of differentiated hepatocytes, these alterations led us to explore the impact of both ß-catenin functions on cell differentiation.

### Alteration of the hepatocyte differentiation and polarity upon ß-catenin knock-downs

Differentiated hepatocytes are characterized by the expression of specific markers, including xenobiotic-metabolizing enzymes, transporters, transcription factors and bile canaliculi molecules. Interestingly, we observed an alteration in mirror upon silencing of either the WT or the mutated allele of ß-catenin. Hepatocyte markers were found up-regulated upon mutated ß-catenin silencing and down-regulated upon WT ß-catenin silencing (Fig. 3A). As HNF4a (hepatocyte nuclear factor-4 alpha) is a transcription factor involved in hepatocyte differentiation, we further explored the impact of ß-catenin KD on HNF4a signaling. We first analyzed the expression of HNF4a by qRT-PCR and found a slight decrease of its expression upon WT ß-catenin KD, whereas it is not significantly altered upon silencing of the oncogenic ß-catenin (Fig. 3B). We found a significant increase of HNF4a transcriptional activity upon silencing of the oncogenic ß-catenin using ApoC3 promoter reporter assay (Fig. 3C). Accordingly, in our transcriptional analysis, expressions of transcriptional positive targets of HNF4a were found upregulated upon silencing of the mutated ß-catenin (Fig. S3 A-C) confirming the results obtained for ApoC3 and ApoM mRNA expressions by qRT-PCR (Fig. 3D). In both assays, the removal of WT ß-catenin slightly decreased the expression of HNF4a target genes. As described earlier^16^, this result confirms that transcriptional activity of ß-catenin may repress the hepatocyte differentiation program of HNF4a. Altogether, these results suggest that the structural function of ß-catenin, mainly supported by the WT ß-catenin, is necessary to maintain a differentiated state of hepatocytes, and that the inhibition of the transcriptional activity of oncogenic ß-catenin reverses the dedifferentiation program of HepG2 cells.

**Figure 3:**
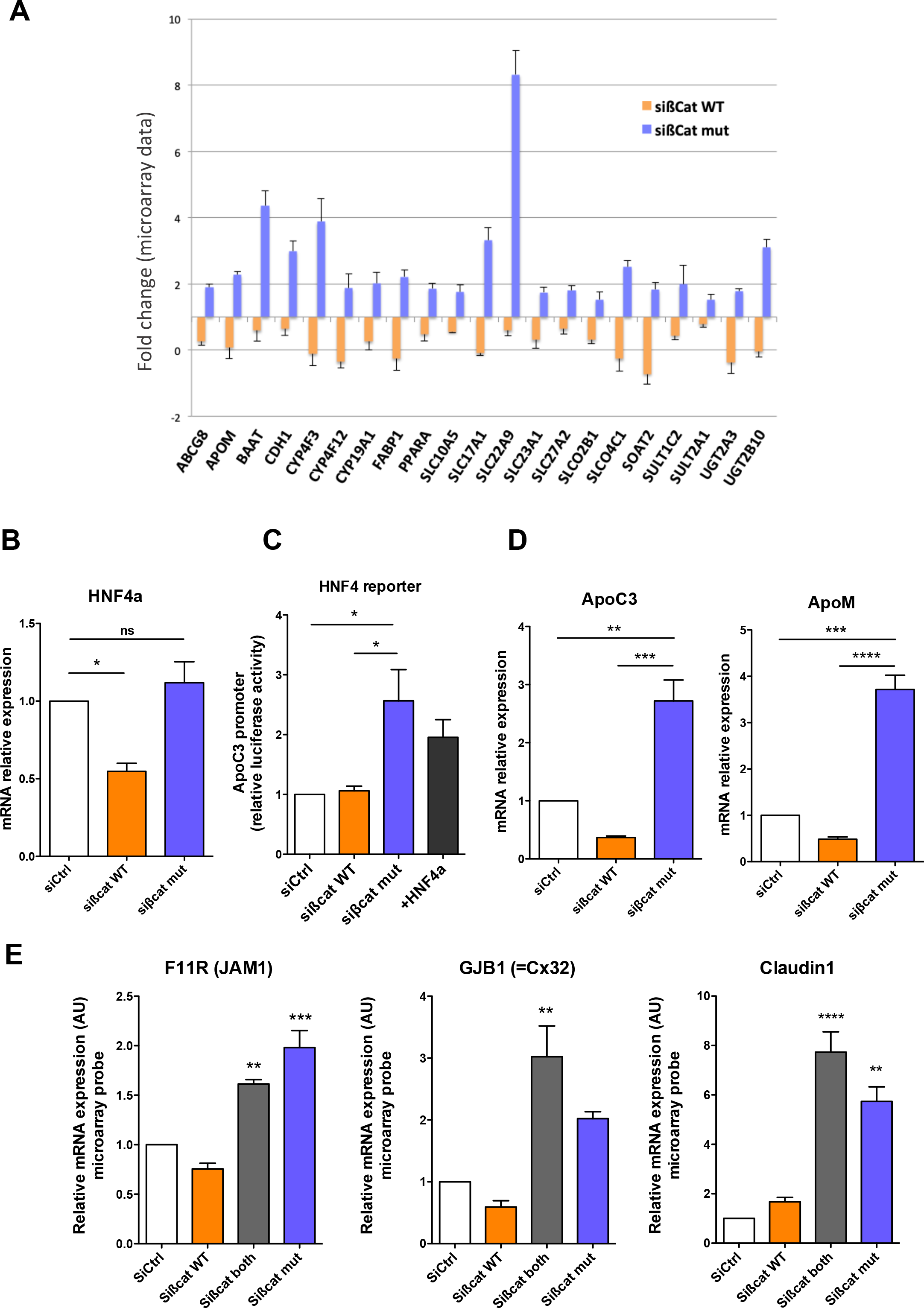
(A) Alteration of the expression of differentiated hepatocyte markers upon silencing of both alleles of ß-catenin in HepG2 cells. The graph shows the relative expression of the indicated genes extracted from the transcriptomic analysis. (B-D) HNF4a activity is up-regulated upon silencing of mutated ß-catenin. (B) Indicated siRNA were transfected in HepG2 cells, and *HNF4A* relative mRNA level was analyzed. Shown is the relative mRNA level compared to control transfected cells. (C) HepG2 cells transfected with indicated siRNAs were transfected with HNF4a responsive luciferase reporter. Shown is the mean relative luciferase activity, normalized to Renilla luciferase and compared to siRNA control transfected cells. (D) Relative mRNA levels of *APOC3* and *APOM*, both positive transcriptional targets of HNF4a were analyzed by qRT-PCR. Shown is the relative mRNA level compared to control transfected cells. (E) Alteration of the expression of polarity markers upon silencing of both alleles of ß-catenin in HepG2 cells. The graph shows the relative expression of the indicated genes extracted from the transcriptomic analysis. (B-E) Each graph shows the quantification of three independent experiments. Error bars indicate s.e.m (n=3), *P* values from one way ANOVA.

As differentiated hepatocytes are polarized epithelial cells endowed with the capacity to produce and excrete bile into a specialized structure called bile canaliculus (BC), we also analyzed the impact of ß-catenin KD on BC markers. As described above for hepatocyte markers, we observed an alteration in mirror in the expression of junction- and polarity-associated genes (Fig. S3D-E), such as JAM-A (*F11R* gene), connexin-32 (*GJB1* gene) and claudin-1 (*CLDN1* gene) (Fig. 3E, S3D-E). As HepG2 cells retained the ability to form BC in culture, we addressed the impact of ß-catenin on their maintenance using confocal microscopy by staining F-actin and radixin (Fig. 4A). Interestingly, we found that depletion of WT β-catenin induced a decrease, whereas depletion of mutated β-catenin an increase, in the number of cells exhibiting BC (Fig. 4A-B). We also performed super-resolution STED microscopy on HepG2 cells stained with phalloidin in order to visualize BC with an improved resolution than confocal (Fig. 4C and Fig. S4A). Quantification of BC features demonstrated that mutated ß-catenin KD increases their size, as attested by the increase of their area, perimeter and diameter, whereas their circularity remains unchanged (Fig. 4D), and the number of cells engaged in the formation of each canaliculus was found higher (Fig. S4B). Finally, to check the functionality of BC, we examined their ability to translocate CFDA, a fluorescent substrate of apical plasma membrane ABC transporters, into the apical lumen. As shown between adjacent HepG2 cells on Figure 4E, removing the mutated form of ß-catenin increased the percentage of CFDA positive BC, demonstrating they are fully functional (Fig. 4E). Our results demonstrated that in tumor hepatocytes, WT and mutated ß-catenin act antagonistically on hepatocyte BC. These results suggest that WT ß-catenin still acts as a gatekeeper of hepatocyte differentiation and polarity, even in tumor hepatocytes.

**Figure 4:**
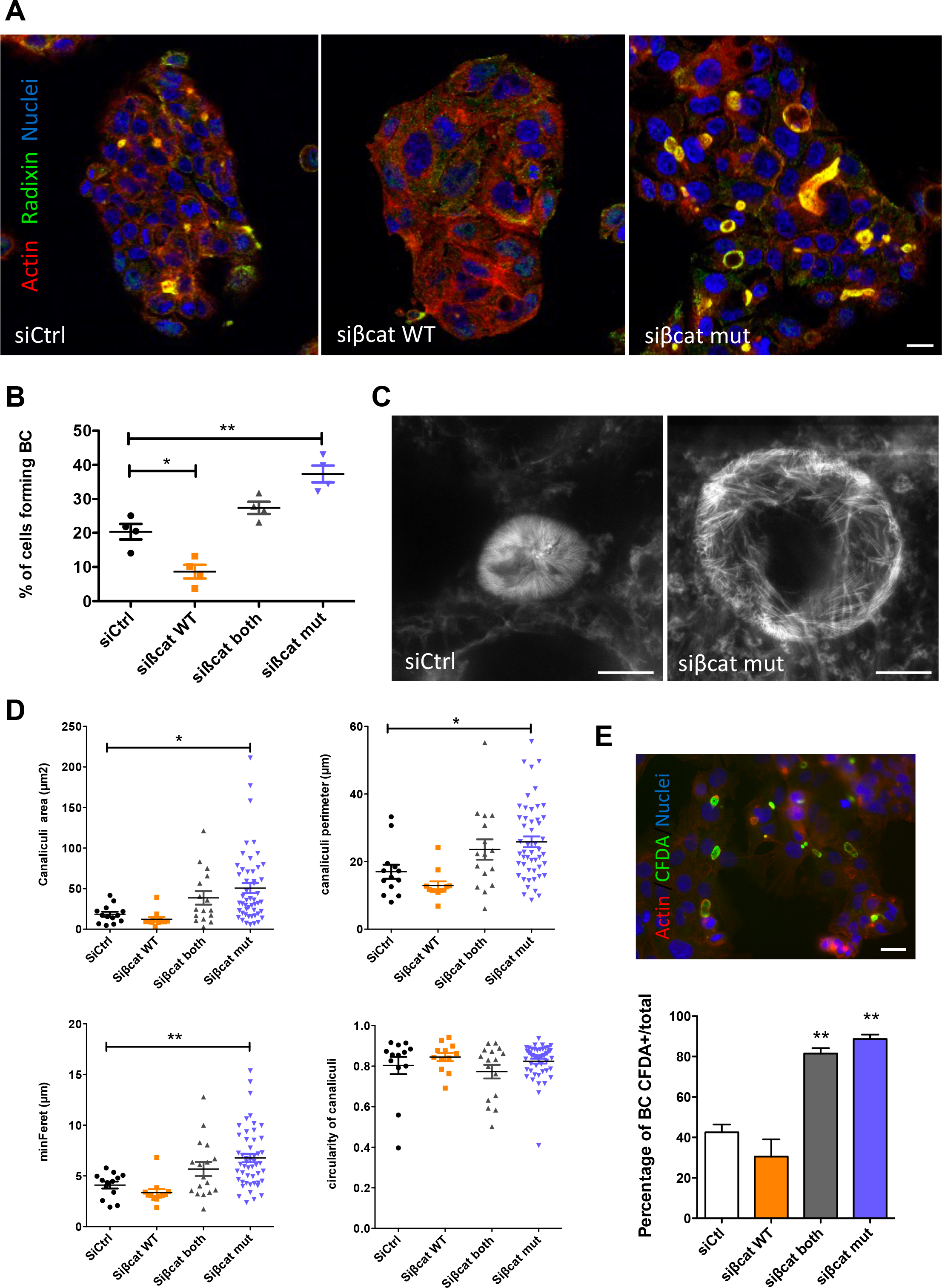
Alteration of bile canaliculi formation upon silencing of ß-catenin in HepG2 cells. (A) HepG2 cells transfected with indicated siRNAs were fixed and stained with phalloidin (red), anti-radixin antibodies (green) and Hoescht (blue). Scale bar: 15 µm. (B) Quantification of the percentage of cells forming BC in the conditions described in (A). Graph shows the quantification of four independent experiments, where at least 100 cells were observed per experiment. (C) siRNA transfected HepG2 cells were fixed, stained with phalloidin-ATTO and observed by STED microscopy. Scale bar: 5 µm. (D) BC features (area, perimeter, circularity, min Feret (diameter)) were quantified by imageJ on STED images performed as described in (C). Each dot corresponds to one BC. (E) Quantification of BC functionality by using CFDA incorporation. Life-Act (red) expressing HepG2 cells were treated with CFDA (green) and observed with fluorescence microscope. Scale bar: 25 µm. Graph shows the quantification of three independent experiments. (B, D-E) Error bars indicate s.e.m (n=4 for B, n=3 for D-E), *P* value from one way ANOVA.

### Fascin-1, as a target of ß-catenin, involved in hepatocyte dedifferentiation

We searched, in our transcriptomic data, for genes antagonistically expressed upon silencing of WT and mutated ß-catenin in HepG2 cells, potentially impacting both their differentiation and polarity. We focused on *FSCN-1* found up-regulated upon WT ß-catenin KD and down-regulated upon mutated ß-catenin KD (Fig. S5A). *FSCN-1* encodes fascin-1, an actin-bundling protein, which is normally not expressed in epithelial cells, i.e. absent in mature hepatocytes. Instead, villin, encoded by the *VIL1* gene, is the actin-bundling protein associated to BC microvilli in differentiated cells.

As villin and fascin-1 that share similar function, were found altered in an opposite manner upon ß-catenin KD (Fig. S3D-E and S5A), we further studied the role of fascin-1 in hepatocyte dedifferentiation. Moreover, fascin-1 is expressed in tumors including HCC^17^ and described as a transcriptional target of β-catenin in breast, gastric and colon cancer cells^18–20^. We first analyzed fascin-1 protein expression in HB and HCC cell lines, with different ß-catenin status (Fig. S5B). We found that fascin-1 was more expressed in cell lines bearing-*CTNNB1* deletion (HepG2) or mutations (SNU398 and Huh6), compared to non-mutated (Huh7, Hep3B) cell lines. The level of Fascin-1 protein was positively correlated to ß-catenin protein level, which accumulates upon mutations (Fig. S5C). As a ß-catenin transcriptional target, fascin-1 mRNA expression was strongly inhibited upon treatment of HepG2 cells with siRNAs sißcat-mut and sißcat-both (Fig. 5A). As previously observed for Axin2 and CyclinD1 (Fig. 1E), the expression of fascin-1 is slightly but significantly upregulated upon removal of WT ß-catenin. This alteration was also detected at the protein level (Fig. S5D). Using a luciferase assay with fascin-1 promoter, we show that this regulation occurs at the transcriptional level, with an antagonistically regulation of promoter activity upon silencing of either WT or mutated ß-catenin (Fig. 5B). Thus, as previously shown in other cancer cells, our results confirm that *FSCN1* is a target gene of β-catenin in tumor hepatocytes. We next wonder whether the variation of fascin-1 expression may play a role in the alteration of hepatocyte differentiation in a tumor context. Interestingly we found that fascin-1 silencing led to an increase of ApoC3, E-cadherin and claudin1 mRNA expressions in HepG2 cells (Fig. 5C). The same tendency was observed in HCC cell lines (Fig. S6A-B). We also found that fascin-1 KD (Fig. 5D) led to an increase of two to three folds in the number of BC in HepG2 cells (Fig. 5E). Moreover, we show that the depletion of fascin-1 upon inhibition of WT β-catenin expression restores the formation of BC (Fig. 5F), demonstrating that fascin-1 is one of the effectors responsible of the impact of ß-catenin on BC formation. Finally, overexpression of fascin-1 induced a decrease in BC formation in comparison to control (Fig. 5G). Altogether these results demonstrated that the level of fascin-1 regulates hepatocyte differentiation status. Moreover, the impact of WT and mutated ß-catenin silencing on hepatocyte differentiation is in part due to fascin-1 expression.

**Figure 5:**
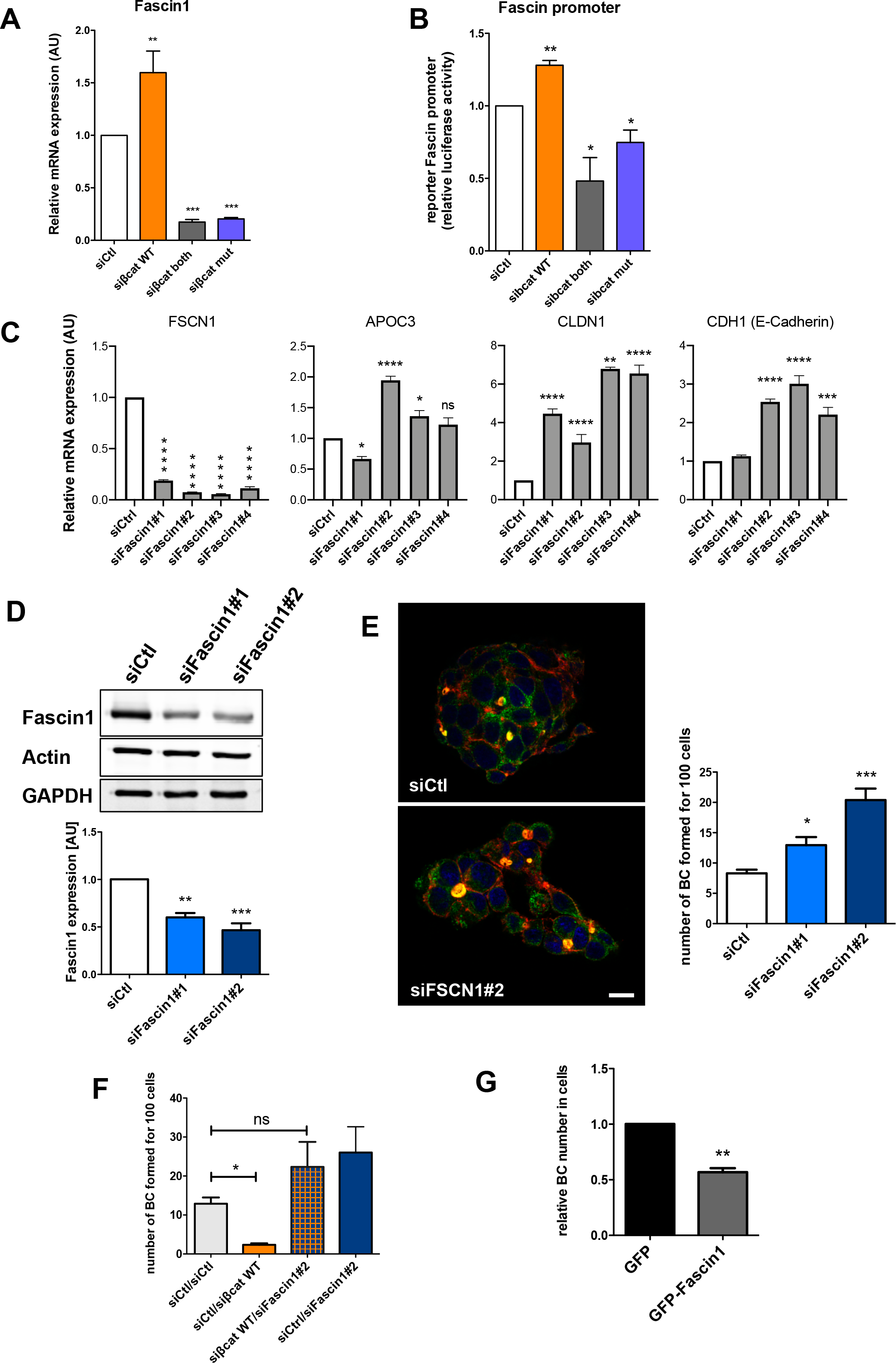
Fascin-1, a target of β-catenin, alters hepatocyte differentiation status. (A) Fascin-1 mRNA expression upon depletion of WT and/or mutated β-catenin analyzed by RT-qPCR in HepG2 cells. (B) Activity of Fascin-1 promoter studied by reporter luciferase assay. Shown is the mean relative luciferase activity, normalized to Renilla luciferase and compared to control siRNA transfected cells. (C) mRNA levels of *FSCN1*, *APOC3*, *CLDN1* and *CDH1* upon treatment of HepG2 cells with siRNA targeting Fascin-1 (siFascin1#1, #2, #3 and #4). (D) HepG2 were transfected with indicated siRNAs, and protein extracts were analyzed using anti-Fascin-1, ß-actin and GAPDH antibodies. The graph shows the quantification of three independent experiments. (E) siRNA transfected HepG2 cells were fixed and stained with phalloidin (red), anti-radixin antibodies (green) and Hoescht (blue). Scale bar: 15 µm. The graph shows the quantification of three independent experiments where the number of BC formed for 100 cells are indicated. (F-G) Experiments were performed as described in (E) with either co-transfection of indicated siRNAs (F) or transfection of pEGFP-C1 (GFP) or pEGFP-C1-Fascin-1 (GFP-Fascin-1) (F). (A-G) Graphs show the quantification of at least three independent experiments. Error bars indicate s.e.m (n=4 for A, n=5 for B and n=3 for C-F), *P* values from one way ANOVA.

### Fascin expression is high in undifferentiated tumors in mice

We next aimed at validating fascin-1 as a ß-catenin target in liver tumors in mice. We used mouse models that mimic ß-catenin dependent tumorigenesis, such as the APC loss-of-function and the Δexon3 ß-catenin models. These models were shown to lead to ß-catenin pathway activation and the development of phenotypically undistinguishable liver tumors within about 10 months^15, 21^. Interestingly, two phenotypically distinct tumors defined as differentiated and undifferentiated tumors are generated in these mice^15, 21^. Well-differentiated tumors are characterized by hepatocyte-like tumor cells that maintain a β-catenin-induced expression of glutamine synthetase (GS) and present nuclear β-catenin. Undifferentiated tumors are composed of small cells with basophilic nuclei, that they strongly express nuclear β-catenin^15^. A RNA-Seq analysis showed that these tumors lose the expression of GS, which is reminiscent of a loss of differentiation (Fig. S7A). We thus made use of this mouse cohort to analyze fascin-1 expression (Table S3). RNA sequencing data from these murine tumors demonstrated that fascin-1 expression is increased in both, well-differentiated and undifferentiated types of tumors in comparison to normal liver (Fig. 6A and S7B). We further highlighted that fascin-1 expression is higher in undifferentiated samples than in the differentiated ones (Fig. 6B). Moreover, fascin mRNA expression was found to correlate positively with various markers of undifferentiated tumors such as *MMP2*, *VIM*, *HIF1A* and *YAP1*, and negatively with markers of differentiated tumors such as *HNF4a*, *APOC3* and *GJB1* (Fig. S7C). Finally, as a bona fide ß-catenin target, fascin-1 expression correlated positively with level of *CTNNB1*, *LEF1* and *TCF4* (Fig. S7C). These new findings were confirmed by immunohistochemistry performed on both types of murine tumors obtained from a new cohort of APC KO mice. As described previously^15^, activation of the β-catenin pathway in the tumor cells was characterized by nuclear localization of the β-catenin along with high expression of GS in well-differentiated tumors (Fig. 6C) and a lower to almost absent expression in undifferentiated ones (Fig. 6D). Fascin-1 staining, which is absent of the non-tumoral hepatocytes, was found increased in both types of tumors. Moreover, whereas its expression is low in differentiated tumors, fascin expression appears high in undifferentiated tumor cells, with a cytoplasmic and membranous staining. Thus, in mice, fascin-1 expression is a marker of ß-catenin-induced undifferentiated tumors. As those murine tumors were found to be transcriptionally close to human mesenchymal HBs^15^, this prompted us to explore Fascin-1 expression in human HBs.

**Figure 6:**
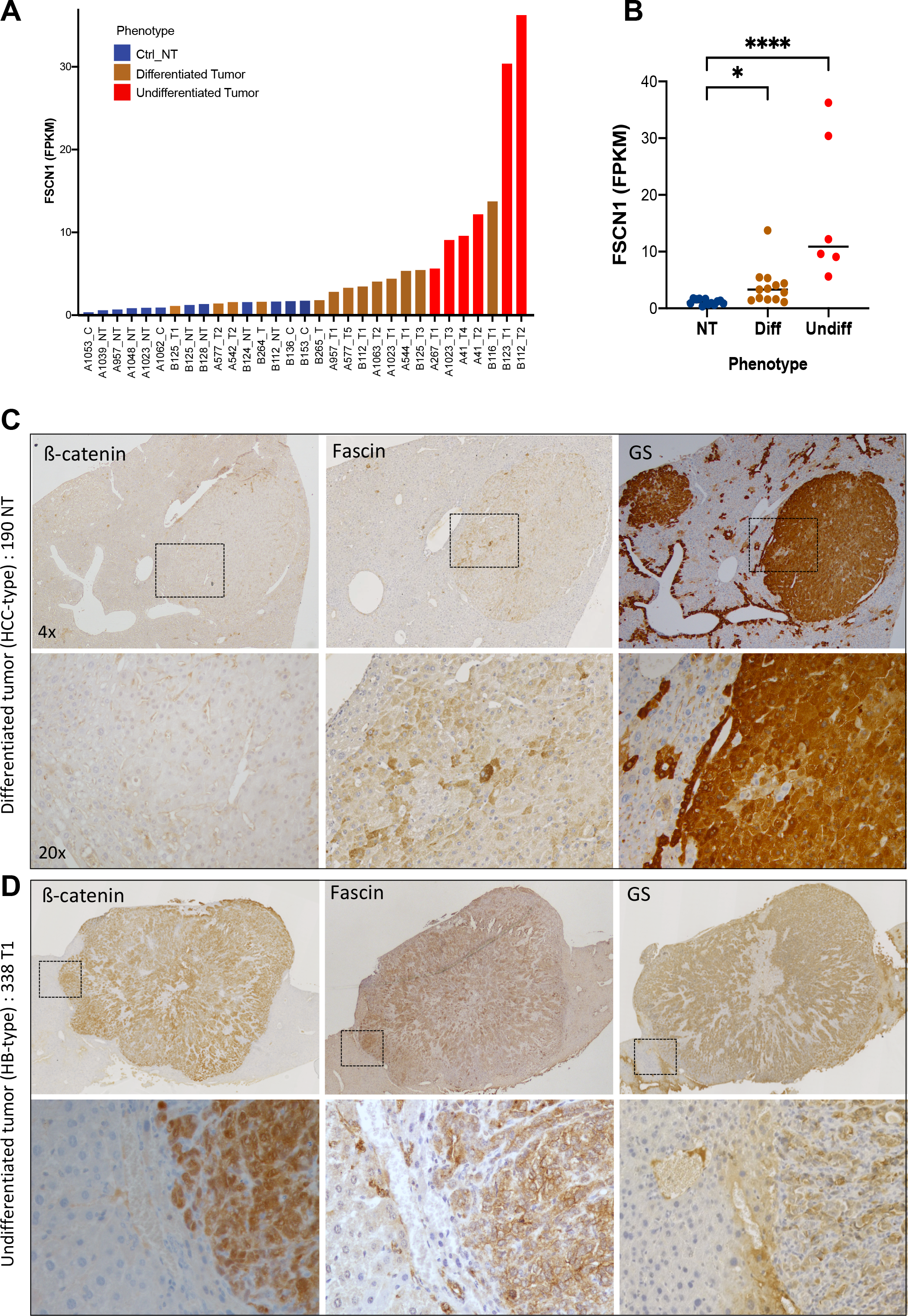
Fascin-1 expression in murine ß-catenin-mediated tumors. (A-B) Data were extracted from RNAseq performed on mouse hepatic tumors induced either from APC KO or ß-catenin Δexon3 expression in livers. *FSCN-1* expression is shown in non-tumoral samples (n=12) and in differentiated (n=13) and undifferentiated (n=6) tumors. *P* values from one way ANOVA. (C) Representative images of immunohistochemistry of ß-catenin, Fascin-1, and glutamine synthetase (GS) in differentiated (HCC-type) (C), and undifferentiated (HB-type) (D) murine tumors. Boxed regions are enlarged in the zoom images.

### Fascin-1 is a marker of embryonal contingent in human HB

In order to analyze fascin-1 expression in human HBs we first made use of public datasets^3, 7^. We found that fascin-1 mRNA is specifically expressed in the C2-subtype of HBs that corresponds to poor-prognosis tumors, with no apparent difference between C2A and C2B subtypes (Fig. 7A). As observed in mice, we found that *FSCN1* mRNA expression correlates negatively with markers of differentiated hepatocytes such as *HNF4a*, *APOC3*, *GJB1* and *CLDN1* (Fig. S8A). We then used immunohistochemistry to confirm expression of the protein product in HB samples. Whereas fascin1 is not expressed in human normal hepatocytes and restricted to sinusoidal cells in normal and peri-tumoral tissues (Fig. S8B), we found that fascin-1 is expressed in a specific contingent of tumor cells in HBs (Fig. 7B). Fascin-1 staining is highly consistent with the results obtained in mice, showing that fascin-1 is expressed in GS-negative and ß-catenin highly positive cells (Fig. 7B and S7C). These cells correspond to the embryonal contingent of undifferentiated small cells with basophilic nuclei. Thus, in human, as found in mice, fascin-1 expression is a marker specific of ß-catenin-induced undifferentiated tumors.

**Figure 7:**
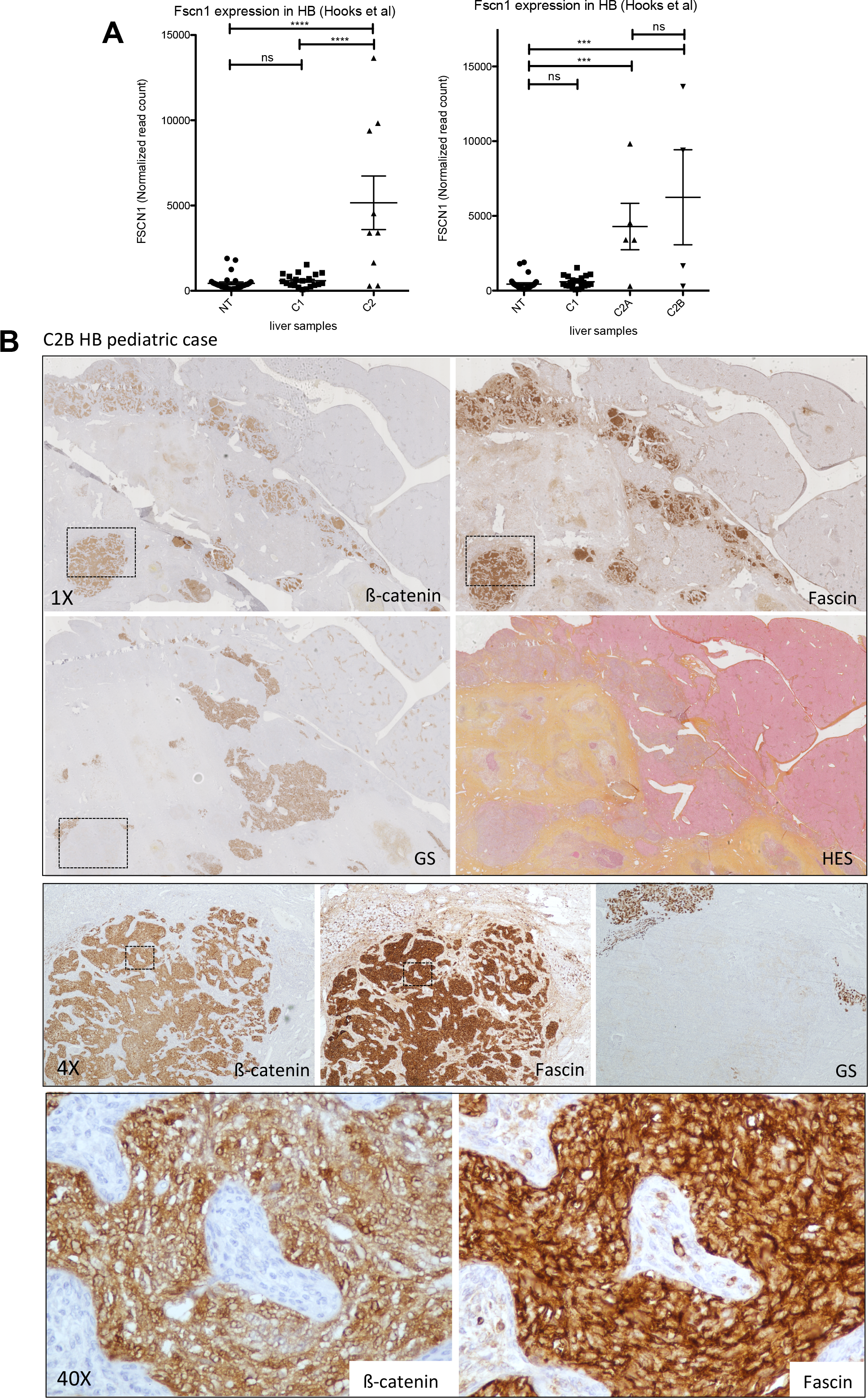
Fascin-1 expression in human HBs. (A) Data were extracted from RNAseq performed on human HBs by Hooks et al.^7^. FSCN-1 expression is shown in non-tumoral samples (n=30) and in C1 (n=20) and C2A or C2B (n=9) tumors. *P* values from one way ANOVA. (B) Representative images of immunohistochemistry of ß-catenin, Fascin-1 and GS and HES staining in a C2B HB pediatric case. Boxed regions are enlarged in the zoom images.

## DISCUSSION

The two functions of ß-catenin, structural at cell-cell junctions and transcriptional in the nucleus are difficult to dissociate and to study individually. Indeed as carried by the same protein, both functions are intrinsically linked. Here, we developed the dual ß-catenin KD HepG2 model, which allow us to address the interplay between the WT and the mutated ß-catenin in liver tumor cells. We found that the mutated form of ß-catenin is dedicated to the transcriptional function, whereas the WT ß-catenin, without Wnt stimulation, is more endowed with its adhesive function in HepG2 cells. Thus, this model is suitable to address independently the structural and the transcriptional functions of ß-catenin. Interestingly, the transcriptomic analysis of the dual ß-catenin KD HepG2 model revealed that disrupting each of the function alter gene expression. Consistent with the transcriptional activity of ß-catenin, genes that were dysregulated upon removal of mutated ß-catenin were coherent with a Wnt/ß-catenin signature. In contrast, genes altered upon silencing of the WT allele were endowed with different signaling pathways that remain to be explored. We found that transcriptional and adhesive activities of β-catenin play antagonistic role in tumor hepatocytes. Whereas the structural function of ß-catenin is necessary to maintain a differentiated state of hepatocytes, the transcriptional activity of ß-catenin induces the dedifferentiation program of HepG2 cells. KD of each allele specifically reverts those programs. As published earlier^16^, we found that this antagonism is in part due to the regulation of the transcriptional activity of HNF4a, required to maintain a hepatocyte differentiation program. Consistent with this antagonism between the transcriptional and the structural activities of ß-catenin we found that WT and mutated ß-catenins act in an opposite manner on hepatocyte polarity.

Hepatocytes polarization involves the formation of functionally distinct sinusoidal (basolateral) and bile canalicular (apical) plasma membrane domains that are separated by tight junctions. A normal membrane polarity is essential for hepatocyte function and its loss may lead to many diseases including cholestasis-associated diseases. A link between β-catenin and BC abnormalities is known since the middle of 2000’s but it remains puzzling. Indeed, on one hand, cholestasis is a key feature of ß-catenin-mutated HCCs^22^, but on the other hand the liver-specific β-catenin KO mice were shown to also develop intrahepatic cholestasis associated with BC abnormalities and bile secretory defect^23^. These data suggest that loss of ß-catenin as well as excessive activation of ß-catenin may lead to cholestasis in the liver. Whether this phenotype is due to structural or transcriptional activity of ß-catenin is largely unknown. Several studies reported the involvement of cell adhesion molecules in the maintenance of BC. In cultured HepG2 cells or embryonic chicken hepatocytes, E-cadherin was shown to be important for BC lumen extension^24, 25^. However, its liver specific knock-out in mice did not alter hepatocyte polarity and BC formation^26^. More recently, hepatocyte specific KD of α-catenin was shown to alter BC, resulting of tight junction disruption and enlarged lumens^27^. Our approach permits to uncouple the different functions of WT or mutated β-catenin revealing their involvement in BC formation. We found that the two forms of ß-catenin have an antagonist role, mutated ß-catenin playing a repressor role by decreasing the number, the size, and the functionality of BC, and WT ß-catenin being important for their maintenance. As demonstrated for the development of bile ducts^28^, ß-catenin has to be kept at the right level as loss of ß-catenin or ß-catenin overactivation is detrimental for BC, formation and/or stabilization.

Our work further highlights the involvement of fascin-1 downstream of ß-catenin in the regulation of hepatocyte differentiation. Fascin-1 is a protein that links actin filaments to form bundles of actin present in filopodia or invadopodia^29^. However, fascin-1 is normally not present in epithelial cell microvilli. In our experiments, fascin-1 behaves as a transcriptional target of ß-catenin. Links between fascin-1 and ß-catenin were reported earlier in the literature. Indeed fascin-1 has been reported to be a transcriptional target of β-catenin/TCF signaling in colon cancer cells showed by the inhibition of fascin promoter activation using dominant-negative *TCF4 in vitro*^18^. In addition, ß-catenin was shown to bind constitutively to fascin promoter in carcinoma cells^30^. However, this regulation of fascin-1 by ß-catenin seems to be cell type-specific as no regulation was observed in breast cancer cells^20^. On the other way around, fascin-1 was found to induce epithelial-mesenchymal transition of cholangiocarcinoma cells and promote breast cancer stem cell function by regulating Wnt/ß-catenin signaling^31, 32^.

One finding of our study is that fascin-1 expression alters hepatocyte differentiation status. We indeed found that fascin-1 is highly expressed in ß-catenin-mutated undifferentiated tumors both in mice and in human. In both, fascin-1 expression correlates negatively with differentiated hepatocyte markers and positively with mesenchymal markers. In addition, *in vitro*, we demonstrated that fascin-1 levels directly act on the hepatocyte polarity and differentiation status. We found that silencing of fascin-1 allows an upregulation of epithelial markers and an increase of BC formation. How fascin-1, an actin-binding protein, may regulate epithelial and mesenchymal gene expression, remains to be explored. Interestingly, Fascin-1 is present in a large list of proteins found to bind mRNA, suggesting a potential role of fascin in post-translational regulation^33^. But it is also now widely accepted that the modulation of the actin cytoskeleton acts on gene expression through mechanotransduction pathways.

Thus, our data demonstrate that fascin-1 is a new target of ß-catenin in the liver that plays a key role in the modulation of hepatocyte differentiation. We found that Fascin-1 is a new marker of the embryonal contingent in HBs. Even if fascin-1 is enriched in the C2 subclass of HBs, we also found fascin-1 staining in C1 samples, reporting the complexity of these tumors. As Fascin-1 is associated to HBs with bad prognosis, fascin-1 immunostaining may offer new diagnostic/pronostic opportunities. Moreover, fascin-1 may be suitable to consider as a new actionable target in these liver pediatric tumors.

## Abbreviations

BC: bile canaliculus
CFDA: 5-carboxyfluorescein diacetate
HB: hepatoblastoma
HCC: hepatocellular carcinoma
HNF4a: hepatocyte nuclear factor-4 alpha
KD: knock-down
STED: stimulated emission depletion
WT: wild-type

## Acknowledgements

We thank Dr D. Vignjevic (Curie Institute, Paris, France) for Fascin-1 DNA constructs. We thank the plateforme GenomEast of Strasbourg (Strasbourg, France). Microscopy was done in the Bordeaux Imaging Center, a service unit of the CNRS-INSERM and Bordeaux University, member of the national infrastructure France Bio Imaging, with the help of Dr. Philippe Legros for STED analysis. We thank Anne-Aurélie Raymond (Inserm U1053, Bordeaux, France) for her help in transcriptomic analysis.

**Author names in bold designate shared co-first authorship**

## SUPPLEMENTARY MATERIALS & METHODS

### 3D Cell growth assay

Three-dimensional multicellular spheroids were prepared by seeding cells on a non-adherent surface. Briefly, 2000 cells were added per well, in Ultra Low Attachment 96-well tissue culture microplates (COSTAR 7007) and incubated at 37°C in complete culture medium. Spheroid formation was assessed 24 h later and spheroid growth was followed using the Incucyte system (Essen BioSciences). Spheroid area was measured using Image J software.

### Transcriptomic analysis

Transcriptomic analysis was performed at the plateforme GenomEast of Strasbourg (Strasbourg, France). Biotinylated single strand cDNA targets were prepared, starting from 150 ng of total RNA, using the Ambion WT Expression Kit (Cat # 4411974) and the Affymetrix GeneChip® WT Terminal Labeling Kit (Cat # 900671) according to Affymetrix recommendations. Following fragmentation and end-labeling, 3 µg of cDNAs were hybridized for 16 hours at 45°C on GeneChip® Human Gene 2.0 ST arrays (Affymetrix) interrogating 24 838 genes and 11 086 LincRNAs represented by approximately 21 probes spread across the full length of the RNA. The chips were washed and stained in the GeneChip® Fluidics Station 450 (Affymetrix) and scanned with the GeneChip® Scanner 3000 7G (Affymetrix) at a resolution of 0.7 µm. Raw data (.CEL Intensity files) were extracted from the scanned images using the Affymetrix GeneChip® Command Console (AGCC) version 3.2. CEL files were further processed with Affymetrix Expression Console software version 1.1 to calculate probe set signal intensities using Robust Multi-array Average (RMA) algorithms with default settings. The raw data were archived in the public GEO data repository with the GEO accession GSE144107. Differentially expressed (DE) genes were identified using the fold-change rank ordering statistics (FCROS) method^13^, which associates an f-value with genes. Small f-values (near zero) are associated with down-regulated genes, while higher values (near 1) are associated with up-regulated genes.

## SUPPLEMENTARY FIGURE LEGENDS

**Figure S1:**
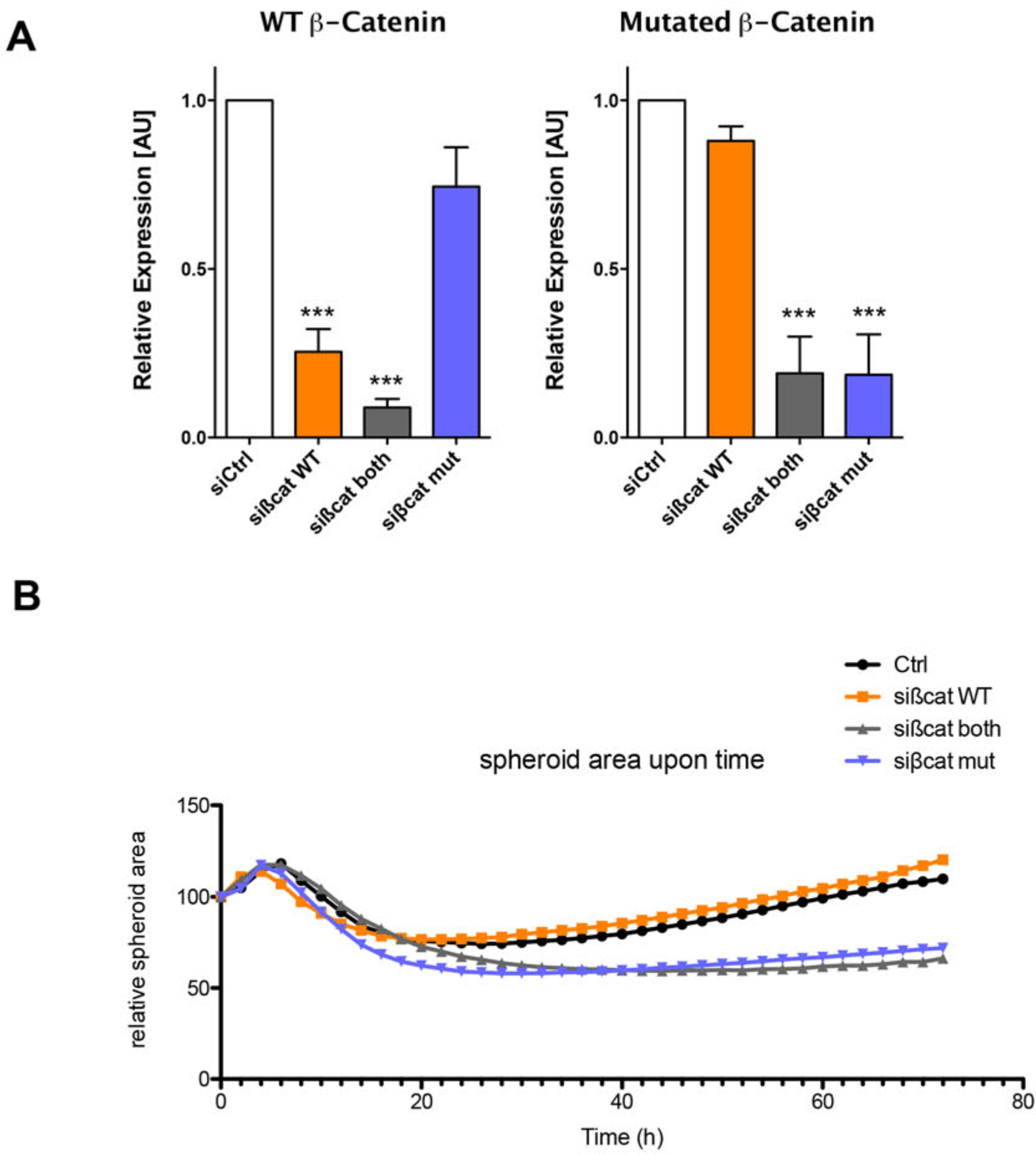
(A) HepG2 were transfected with indicated siRNAs and protein extracts analyzed by immunoblotting using anti-ß-catenin and GAPDH antibodies. Graphs show the quantification of WT (left-hand graph) and mutated (right-hand graph) ß-catenin protein levels normalized to GAPDH. Shown is the relative protein level compared to control transfected cells. Error bars indicate s.e.m (n=3) *** P < 0.001 by one way ANOVA. (B) HepG2 cells transfected with indicated siRNAs were seeded in non-adherent 96-well plates and growth of spheroids was followed using the Incucyte system. The graph shows the mean area of 6 spheroids upon time. This experiment is representative of the three performed.

**Figure S2:**
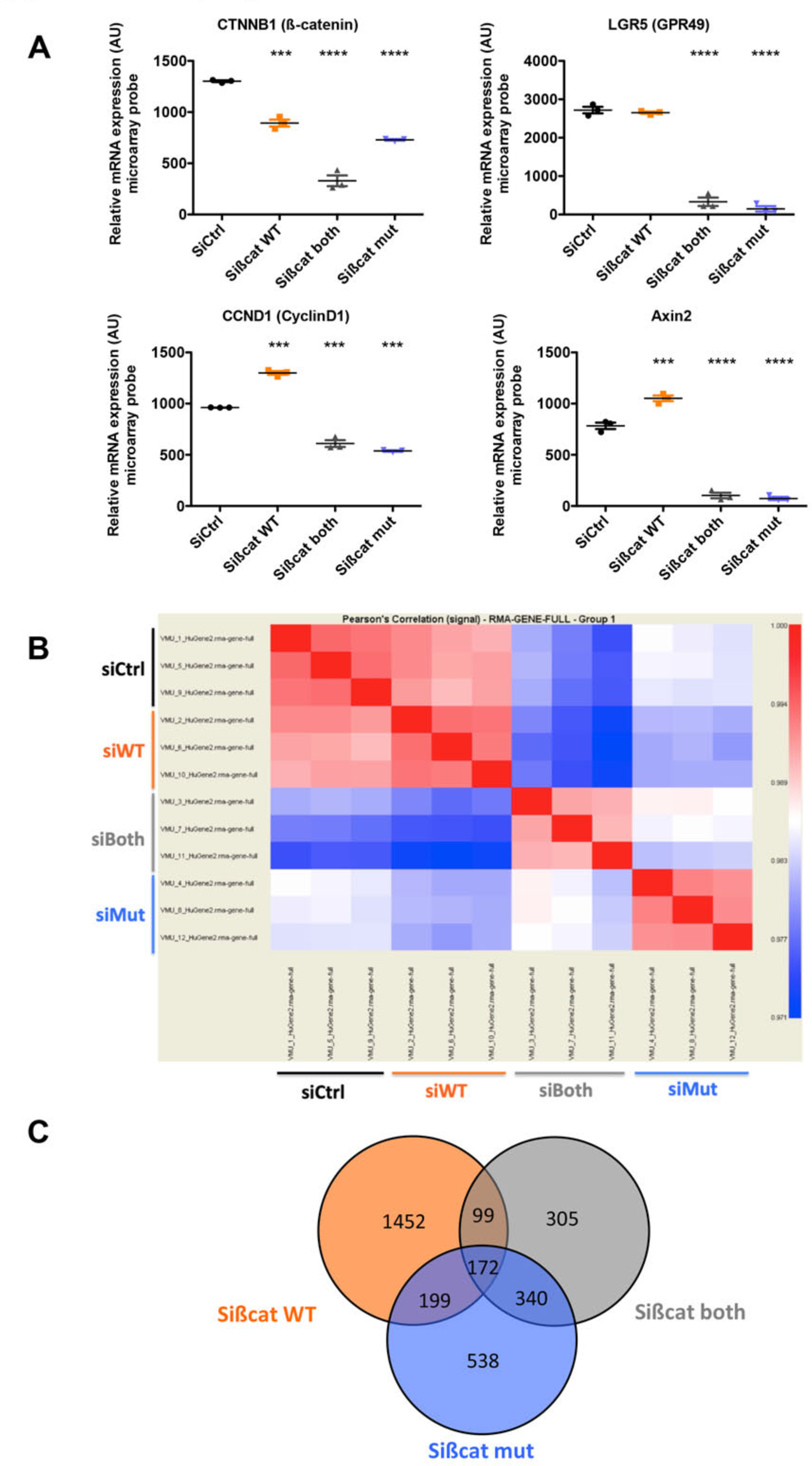
Transcriptomic analysis of the dual ß-catenin KD HepG2 model. (A) Data extracted from the microarray, showing the expression alteration of *CTNNB1*, *LGR5*, *CCND1* and *AXIN2* genes, encoding respectively ß-catenin, GPR49, cyclinD1 and Axin2. Each graph shows the quantification of the three replicates. Error bars indicate s.e.m *** P < 0.001; **** P < 0.0001 by one way ANOVA. (B) Pearson’s correlations between each dataset obtained by Affimetrix. Datasets 1, 5, 9 correspond to HepG2 transfected with control siRNA. Datasets 2, 6, 10 correspond to HepG2 transfected with sißCat WT. Datasets 3, 7, 11 correspond to HepG2 transfected with sißCat both. Datasets 4, 8, 12 correspond to HepG2 transfected with sißCat mut. The color code indicates Pearson’s correlation. (C) Venn diagram of altered genes in sißCat WT, sißCat mut and sißCat both conditions.

**Figure S3:**
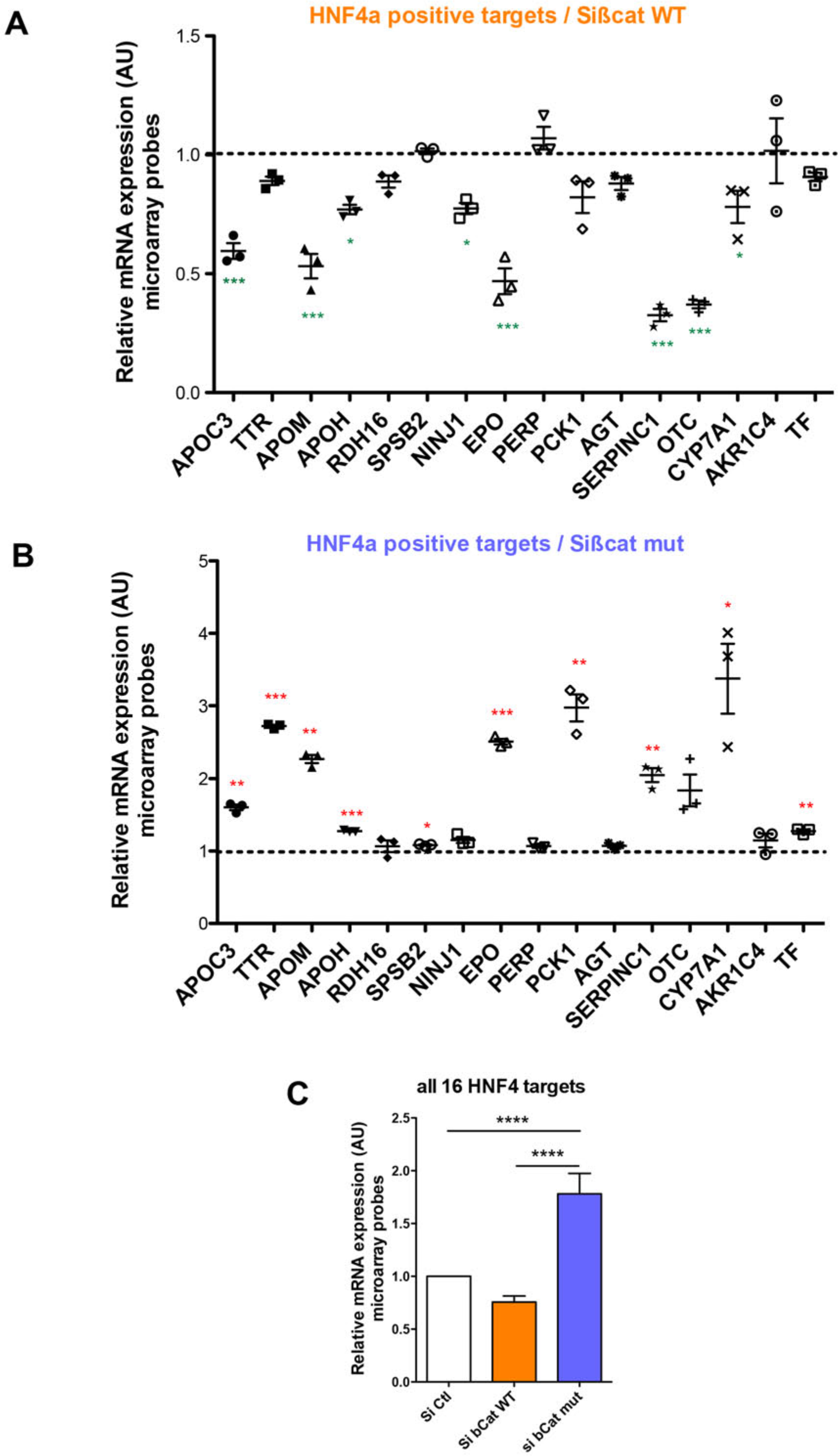
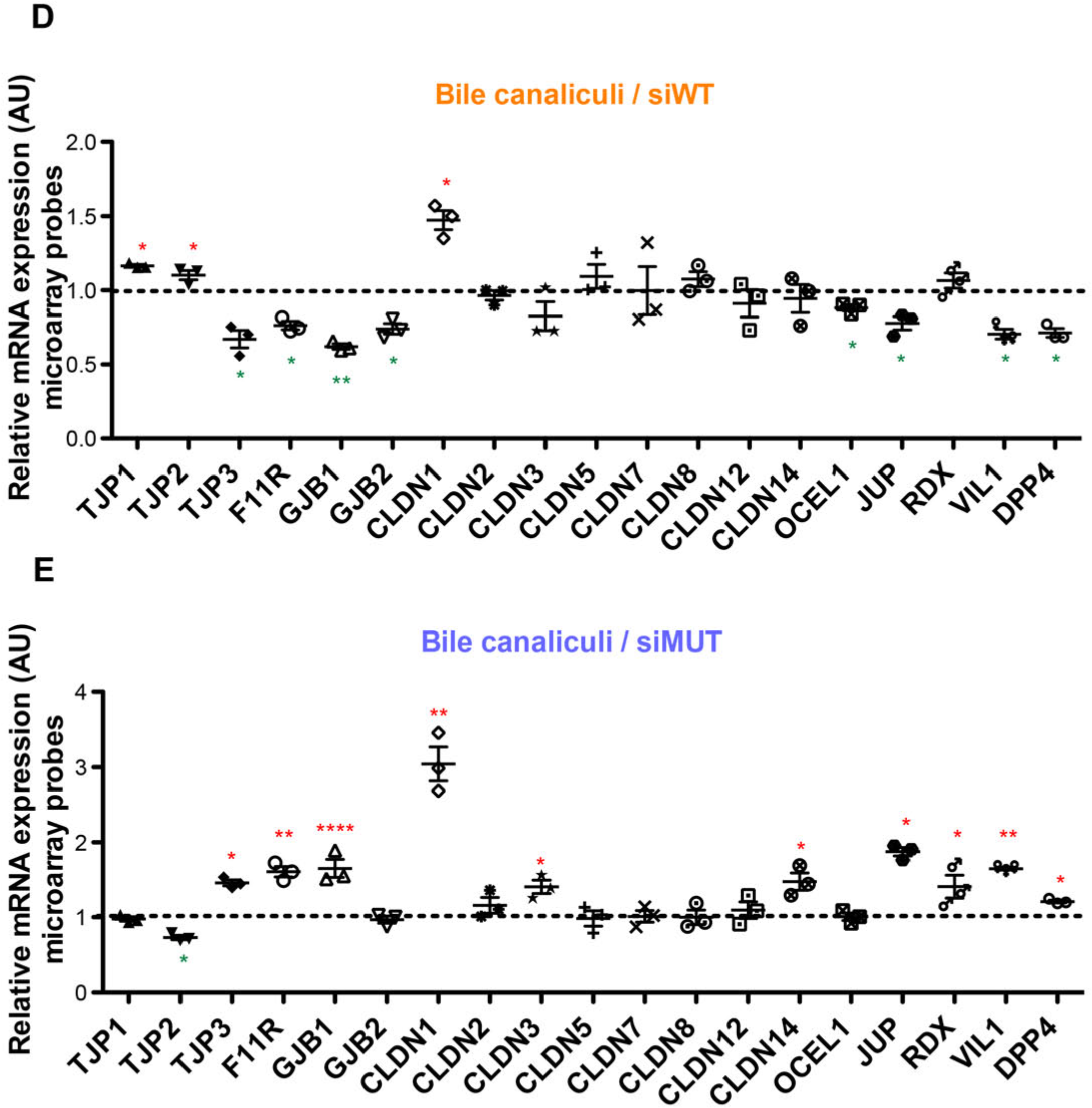
Analysis of the expression of 16 HNF4a positive transcriptional targets extracted from the transcriptomic data, upon silencing of WT ß-catenin (A) or mutated ß-catenin (B) in HepG2 cells. The graphs show the relative expression of the indicated genes extracted from the transcriptomic analysis. Each graph shows the quantification of three independent experiments. Error bars indicate s.e.m (n=3) * P < 0.05; ** P < 0.01; *** P < 0.001 by *t*-test compared to control. (C) Pooled data of the 16 HNF4a targets. **** P < 0.0001. (D-E) Alteration of the expression of polarity-associated genes upon silencing of both alleles of ß-catenin in HepG2 cells. Graphs show the relative expression of the indicated genes extracted from the transcriptomic analysis. Each graph shows the quantification of three independent experiments. Error bars indicate s.e.m (n=3) * P < 0.05; ** P < 0.01; **** P < 0.0001 by *t*-test compared to control.

**Figure S4:**
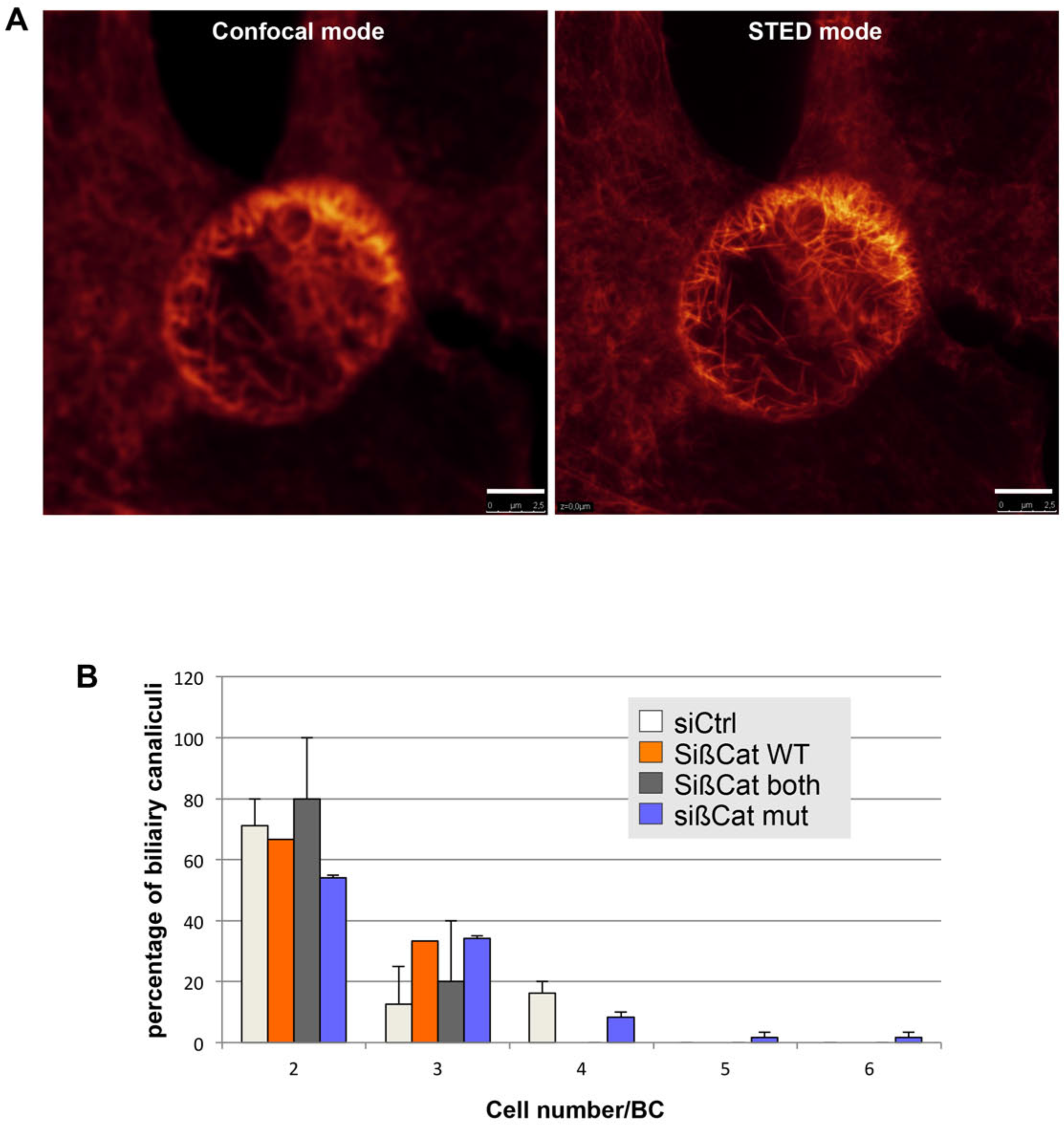
(A) HepG2 cells transfected with siRNA targeting mutated ß-catenin were fixed, stained with phalloidin-ATTO and observed by confocal (left-hand) and STED (right-hand) microscopy. Scale bar: 2.5 µm. (B) HepG2 cells transfected with indicated siRNAs were fixed and stained with phalloidin (red), anti-radixin antibodies (green) and Hoescht (blue). The number of cells engaged in the formation of bile canaliculi was evaluated under the microscope. Graph shows the quantification of three independent experiments. Error bars indicate s.e.m.

**Figure S5:**
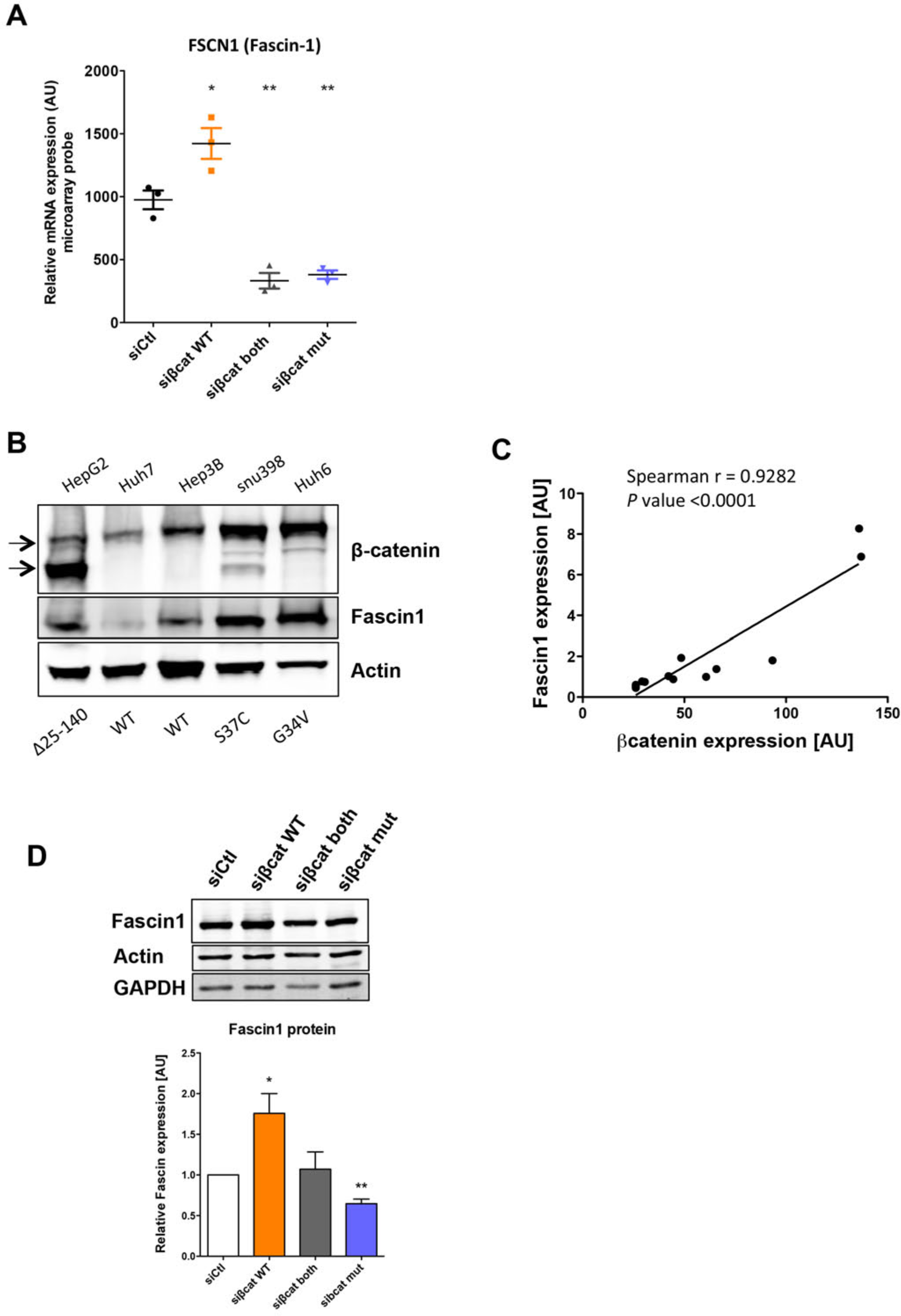
Regulation of Fascin-1 expression by β-catenin. (A) Data extracted from the transcriptomic data, upon silencing of WT ß-catenin and/or mutated ß-catenin in HepG2 cells, showing the expression of *FSCN1* gene, encoding Fascin-1. The graph shows the quantification of the three replicates. Error bars indicate s.e.m. * P < 0.05; ** P < 0.01 by one way ANOVA. (B) Protein extracts from indicated cell lines were analyzed by immunoblotting using anti-ß-catenin, Fascin-1 and ß-actin antibodies. Arrows indicated mutated (about 76 kDa) and WT (about 92 kDa) ß-catenins in HepG2 cells. Status of *CTNNB1* gene in cells is indicated below the immunoblot. (C) Correlation of the level of Fascin-1 and β-catenin in various cell lines, as quantified by immunoblot. (D) Fascin-1 protein expression upon depletion of WT and/or mutated β-catenin observed by western blot. Actin and GAPDH serve as loading controls.

**Figure S6:**
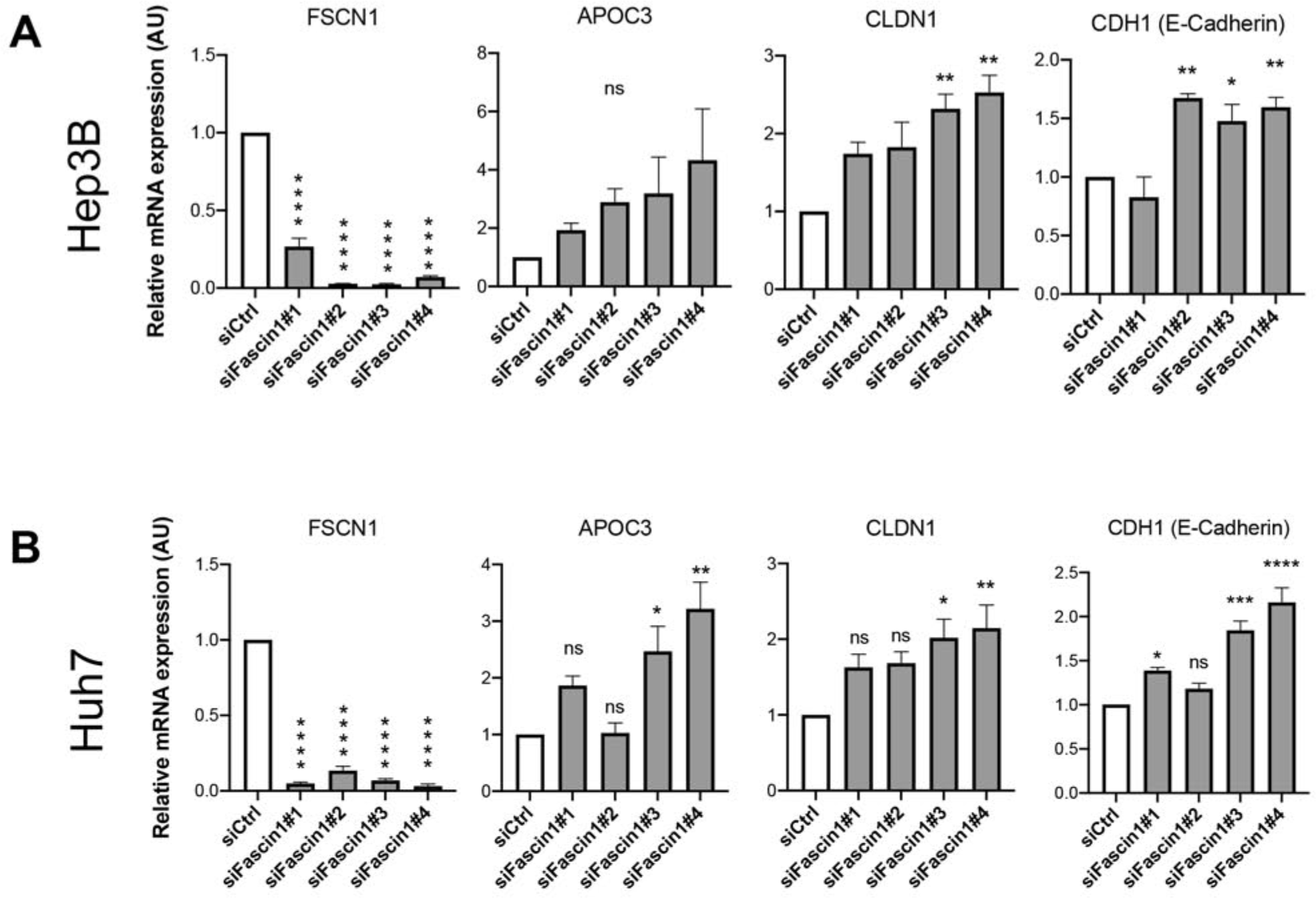
(A-B) mRNA levels of *FSCN1*, *APOC3*, *CLDN1* and *CDH1* upon treatment of Hep3B (A) and Huh7 (B) cells with siRNA targeting Fascin-1 (siFascin1#1, #2, #3 and #4). Graphs show the quantification of three independent experiments. Error bars indicate s.e.m. * P < 0.05; ** P < 0.01; *** P < 0.001; **** P< 0.0001 by one way ANOVA.

**Figure S7:**
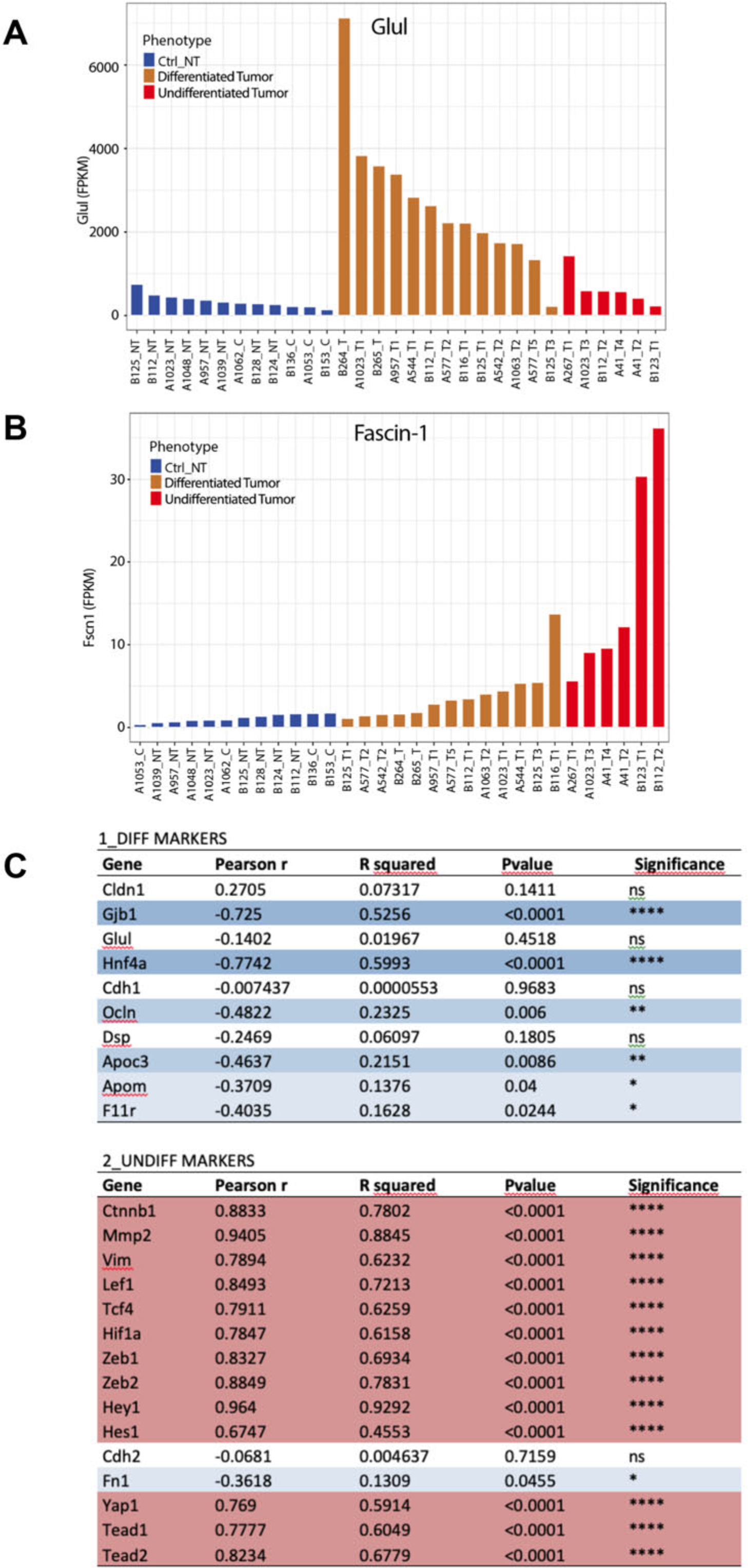
Fascin-1 expression in murine activated ß-catenin-mediated tumors. (A-C) Data were extracted from analyses performed on mouse ß-catenin-mediated tumors by Loesch et al. (Loesch et al., submitted). GLUL encoding glutamine synthetase (A) and FSCN-1, encoding Fascin-1 (B) are shown in non-tumoral samples (n=12) and in differentiated (n=13) and undifferentiated (n=6) tumors. (C) *FSCN1* expression correlates negatively with differentiation markers (highlighted in blue) and positively with undifferentiation markers (highlighted in red) in mouse liver tumors.

**Figure S8:**
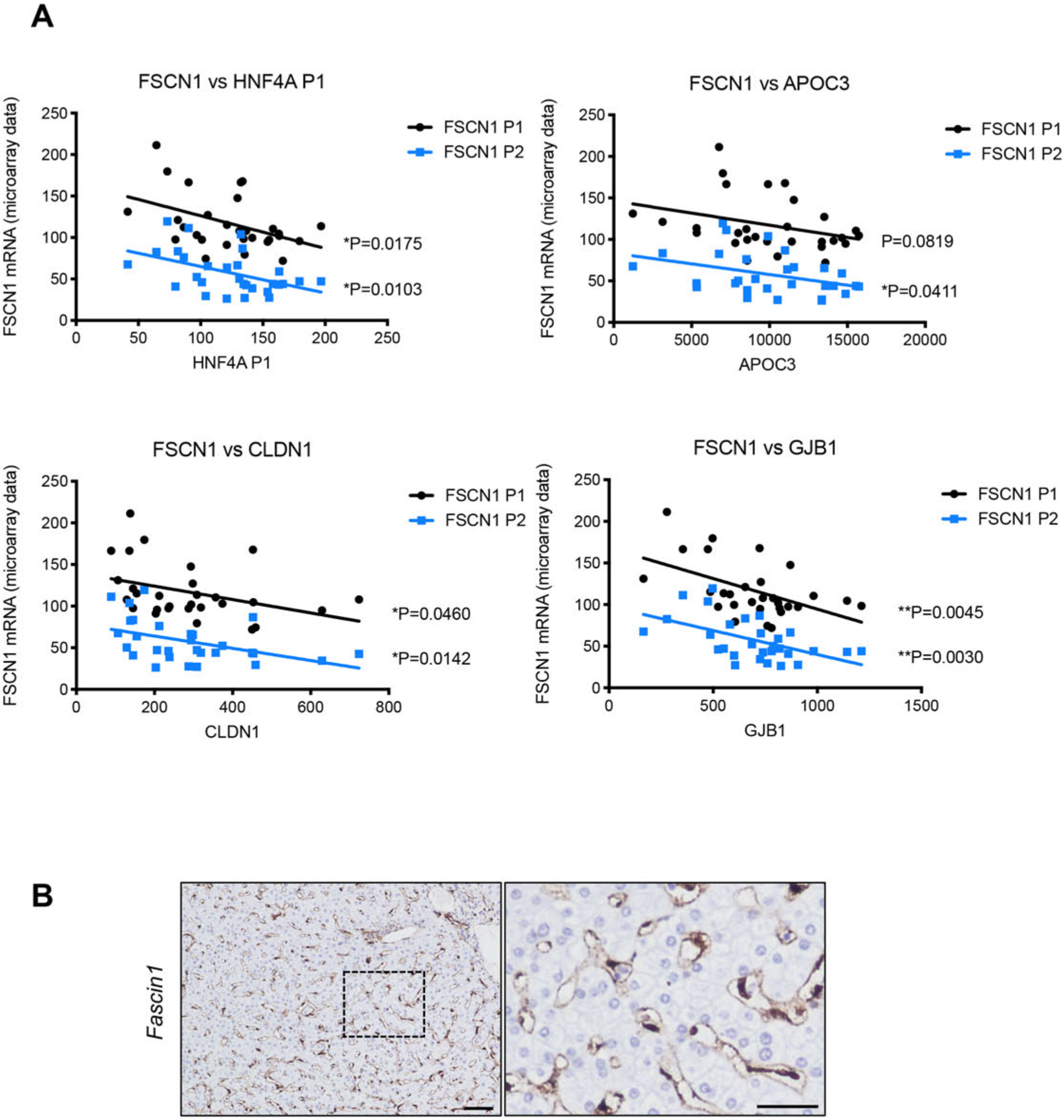
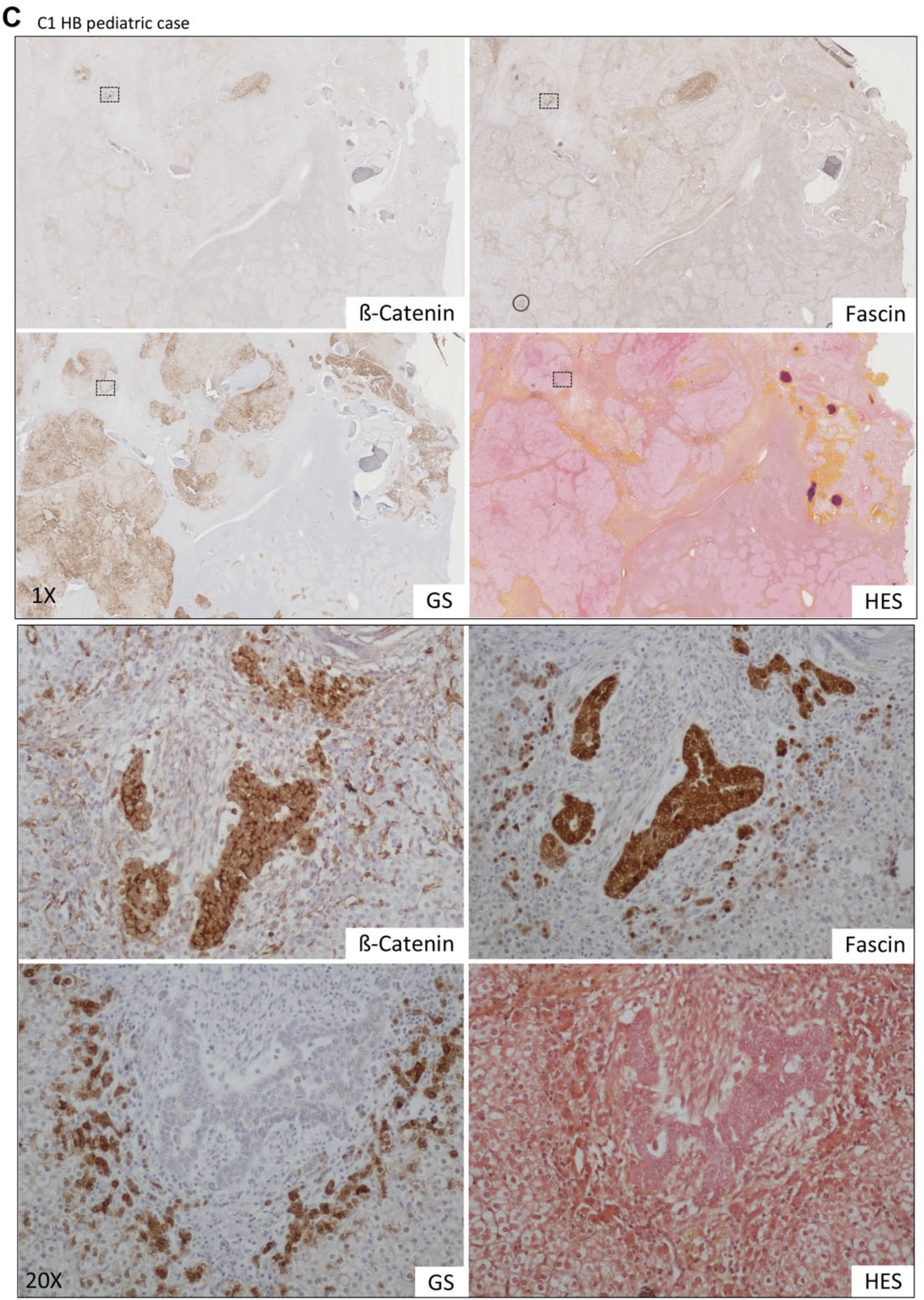
Fascin-1 expression in human HBs. (A) Data were extracted from published analyses from performed on human HBs by Cairo et al., 2008. *FSCN1* expression correlates negatively with *HNF4A*, *APOC3*, *CLDN1* and *GJB1* in human HBs. (B) Representative image of immunohistochemistry of Fascin-1 in a normal human liver. (C) Representative images of immunohistochemistry of ß-catenin, Fascin-1 and glutamine synthetase (GS), and HES staining in a C1 HB pediatric case. Dashed rectangles show part of the images showed in the zoom images.

## Supplementary Tables

<u>Supplementary Table 1:</u> List of the antibodies used in the study.

<u>Supplementary Table 2:</u> List of the qRT-PCR primers used in the study. Forward and reverse primers are indicated for each gene.

<u>Supplementary Table 3:</u> Mouse cohorts used in this study.

<u>Supplementary Table 4:</u> Clinical data of patient samples used for immunohistochemistry analyses.

<u>Supplementary Tables 5-7</u>: List of the deregulated genes upon WT (Table S2), mutated (Table S3) or both (Table S4) ß-Catenin KD in HepG2 cells. Differentially expressed genes were identified using the fold-change rank ordering statistics (FCROS) method (Dembélé D, Kastner P. BMC Bioinformatics 2014;15:14) and which associates an f-value with genes. Small f-values (near zero) are associated with down-regulated genes (highlighted in green), while higher values (near 1) are associated with up-regulated genes (highlighted in pink). “ttest” et “sam” correspond to statistical analyses using, respectively, the Student *t*-test and the Significant Analysis of Microarrays (http://statweb.stanford.edu/~tibs/SAM/) methods.

<u>Supplementary Table 8:</u> Analysis with FuncAssociate 3.0 using the genes listed on tables S5-7.

